# Integrin signaling is critical for myeloid-mediated support of T-cell acute lymphoblastic leukemia

**DOI:** 10.1101/2022.01.05.475106

**Authors:** Aram Lyu, Seo Hee Nam, Ryan S. Humphrey, Tyler A. Durham, Zicheng Hu, Dhivya Arasappan, Terzah M. Horton, Lauren I. R. Ehrlich

**Affiliations:** Department of Molecular Biosciences, The University of Texas at Austin, Austin, TX; Bakar Computational Health Sciences Institute, University of California, San Francisco, San Francisco, CA; Center for Biomedical Research Support, The University of Texas at Austin, Austin, TX; Department of Pediatrics, Baylor College of Medicine/Dan L. Duncan Cancer Center and Texas Children’s Cancer Center, Houston, TX; Department of Oncology, Livestrong Cancer Institutes, The University of Texas at Austin Dell Medical School, Austin, TX

**Keywords:** T-cell acute lymphoblastic leukemia (T-ALL), tumor microenvironment, tumor-associated myeloid cells, cell-cell interactions, integrin signaling

## Abstract

We previously found that T-cell acute lymphoblastic leukemia (T-ALL) requires support from tumor-associated myeloid cells, which activate IGF1R signaling in the leukemic blasts. However, IGF1 is not sufficient to sustain T-ALL survival in vitro, implicating additional myeloid-mediated signals in T-ALL progression. Here, we find that T-ALL cells require close contact with myeloid cells to survive. Transcriptional profiling and in vitro assays demonstrate that integrin-mediated cell adhesion and activation of the downstream FAK/PYK2 kinases are required for myeloid-mediated support of T-ALL cells and promote IGF1R activation. Consistent with these findings, inhibition of integrins or FAK/PYK2 signaling diminishes leukemia burden in multiple organs and confers a survival advantage in a mouse model of T-ALL. Inhibiting integrin-mediated cell adhesion or FAK/PYK2 also diminishes survival of primary patient T-ALL cells co-cultured with myeloid cells. Furthermore, elevated integrin pathway gene signatures correlate significantly with myeloid enrichment and an inferior prognosis in pediatric T-ALL patients.

**Statement of significance:** Although tumor-associated myeloid cells provide critical support for T-ALL, our understanding of the underlying mechanisms remains limited. This study reveals that integrin-mediated adhesion and signaling are key mechanisms by which myeloid cells promote survival and progression of T-ALL blasts in the leukemic microenvironment.

## Introduction

T-cell acute lymphoblastic leukemia (T-ALL) is a hematologic malignancy arising from neoplastic transformation of T-cell progenitors. ALL is the most common pediatric malignancy, with T-ALL accounting for ∼15% and ∼25% of pediatric and adult ALL cases, respectively (1). T-ALL is associated with infiltration of leukemic blasts into multiple organs, including the spleen, bone marrow (BM), lymph nodes (LN), liver, and central nervous system (2). Although current intensified chemotherapy regimens have significantly improved prognoses, with 5-year survival rates for pediatric T-ALL exceeding 90% (3,4), these therapies are toxic and are associated with long-term morbidities, such as neurological toxicity, cognitive impairment, and metabolic disorders (5,6). Moreover, the survival rate for relapsed patients is less than 25% (7–9). Therefore, identifying targets for development of less toxic and more effective therapies would be clinically beneficial. Multiple genetic drivers of T-ALL, including activating mutations in *NOTCH1*, deletion of the *CDKN2A* tumor suppressor locus, and aberrant expression of transcription factors, such as T-cell acute lymphocytic leukemia 1 (TAL1) and LIM domain only 2 (LMO2), have been identified (10–13). Additionally, recent unbiased genome-wide profiling analyses have expanded our understanding of the genetic landscape of T-ALL, identifying novel driver genes and associated molecular mechanisms that promote leukemia initiation and progression (14). However, despite multiple pro-leukemic genomic lesions, T-ALL cells cannot survive in vitro in the absence of cytokines or supportive cell types (15,16), suggesting that cells or signals in the leukemia-microenvironment are required to support leukemia growth and could thus serve as therapeutic targets.

The tumor microenvironment (TME), which is shaped by interactions between tumor cells and their non-transformed neighbors, plays an essential role in promoting the progression of both solid tumors and hematologic malignancies (17,18). Several cell types in the TME have been shown to support T-ALL survival and progression, including vascular endothelial cells through a C-X-C motif chemokine receptor 4 (CXCR4)-CXCL12 axis in the BM (19,20), and thymic epithelial cells (TECs) through release of interleukin 7 (IL-7) and expression of NOTCH1 ligands (11,21,22). We have previously shown that tumor-associated myeloid cells provide critical support for T-ALL, in part by secreting IGF1 to activate IGF1R signaling in leukemic cells (15,16). However, IGF1 is not sufficient to support T-ALL survival in culture, and myeloid cells sensitize T-ALL cells to IGF1 (16), suggesting a role for additional molecular mechanisms underlying myeloid-mediated T-ALL support. We have previously shown that relative to healthy thymocytes, T-ALL cells make frequent and prolonged contact with DCs in the thymus (15), suggesting physical interactions could play a critical role in myeloid-mediated T-ALL support.

Integrins are known mediators of cellular interactions; there are 24 heterodimeric integrin receptors in humans, each consisting of an α and β chain (23). Integrins bind a variety of ligands, including extracellular matrix (ECM) components like fibronectin, collagen, and laminin, as well as adhesion molecules like intercellular adhesion molecule 1 (ICAM-1) and vascular cell adhesion molecule 1 (VCAM-1), expressed by antigen-presenting cells (APCs) and endothelial cells (24). Upon ligand binding, integrins undergo a conformational change, adopting a high-affinity form that recruits cytoskeletal structures via talin and other linker proteins (25). Ligand-bound integrins activate downstream kinases, including focal adhesion kinase (FAK) and proline-rich tyrosine kinase 2 (PYK2), leading to diverse cellular outcomes including cell migration, survival, and proliferation (25). Integrin-mediated interactions with stromal cells or APCs are essential for T-cell development in the thymus and T-cell homeostasis, migration, and activation in the periphery (26–28). Notably, integrins signal cooperatively with growth factor receptors, including IGF1R (25,29,30), and dysregulated integrin signaling has been shown to promote tumorigenesis, metastasis, and chemotherapeutic resistance in a variety of cancers (25,31). Tumor cells can disrupt local ECM and recruit host cells that express integrin ligands, inducing hyperactivated integrin signaling (32,33). Dysregulated integrin signaling has been implicated in tumor progression across multiple malignancies, including breast cancer (34), pancreatic cancer (35), and multiple myeloma (36,37). Integrins have also been implicated in T-ALL pathogenesis, as ICAM-1-dependent interactions with BM stromal cells support T-ALL cell survival in vitro (38), and activation of integrin β1 promotes chemoresistance of T-ALL cell lines (39). Given that integrin signaling promotes survival of healthy T cells and tumors, along with our previous observation that T-ALL cells make frequent and prolonged contact with myeloid cells in the thymic TME (15), we hypothesized that integrin signaling might promote myeloid-mediated support of T-ALL.

Here, we report that survival of primary mouse T-ALL cells is dependent on close contact with tumor-associated myeloid cells. Transcriptional profiling revealed elevated expression of integrins and enrichment of cell-cell adhesion pathways in T-ALL cells, suggesting integrins could promote adhesion to myeloid cells. Consistent with this possibility, inhibiting integrin-mediated cell adhesion between T-ALL cells and myeloid cells results in a significant decrease in T-ALL survival in vitro. Notably, blockade of ICAM-1 and/or VCAM-1 in leukemic mice, or downstream FAK/PYK2 pathways, significantly diminishes T-ALL burden in multiple organs and confers a significant survival benefit in leukemic mice. Consistent with results from mouse models, integrin-mediated cell adhesion and FAK/PYK2 signaling are required for myeloid-mediated support of primary T-ALL cells from pediatric patients. Elevated integrin pathway gene signatures are also associated with an enrichment of myeloid gene signatures and inferior outcomes in pediatric T-ALL patients. Collectively, these results reveal that tumor-associated myeloid cells promote leukemia survival and progression by activating integrin signaling in mouse models of T-ALL and in primary human T-ALL.

## Results

### Close contact is required for myeloid-mediated T-ALL support and integrins are candidate mediators of these interactions

We previously found that close contact between T-ALL cells and leukemia-associated stromal cells promotes T-ALL survival (15), but the importance of physical interactions between T-ALL cells and tumor-associated myeloid cells has yet to be directly tested. To test this, congenic mice were engrafted with primary LN3 T-ALL cells (40). Once mice became leukemic, splenic T-ALL cells were cultured in the presence or absence of tumor-associated myeloid cells in transwell plates, such that the two cell types could either contact one another or were separated by micropores through which only soluble factors could pass (Figure 1A). T-ALL cells survived only when co-cultured in the same chamber with tumor-associated myeloid cells (Figure 1B), indicating cellular contact is required for myeloid cells to support T-ALL survival. Tumor-associated myeloid cells support survival of T-ALL cells, but not healthy T cells (16), suggesting differences in gene expression between healthy T cells and T-ALL cells could impact their ability to receive supportive signals from tumor-associated myeloid cells. Therefore, we re-evaluated our published RNA-seq datasets of mouse primary LN3 T-ALL cells and healthy T-lineage cells from the thymus and spleen (GSE150096) (16). Gene ontology analysis showed that gene signatures of “cell-cell adhesion” were significantly enriched in thymic T-ALL cells relative to healthy thymocytes (*P* = 1.49×10^−9^; Supplementary Figure 1A). Furthermore, the “Integrin1 (Integrin β1; ITGβ1)” pathway is preferentially enriched in T-ALL cells (Supplementary Figure 1B-C), and ITGβ1 transcript and protein levels are elevated in T-ALL cells (Figure 1C-D, Supplementary Figure 1D). Among ITGβ1 heterodimers, ITGα4β1 (VLA-4) and ITGα9β1 mediate cell-cell interactions via binding to VCAM-1 (24). Both the transcript and protein levels of ITGα9, but not ITGα4, are significantly higher in T-ALL cells relative to controls (Figure 1C-D, Supplementary Figure 1E). The “ILK (Integrin-linked kinase)” pathway is also enriched in T-ALL relative to healthy CD8^+^ T cells in the spleen (Supplementary Figure 1F-G), with elevated levels of ILK in T-ALL cells (Supplementary Figure 1H-I). Activation of ILK is associated with engagement of ITGαLβ2 (LFA-1), and LFA-1 promotes T-cell activation upon binding its ligand ICAM-1 on APCs (28,41,42). Thus, we investigated expression of LFA-1 by T-ALL cells and found it is significantly elevated in T-ALL relative to T cells (Figure 1C-D), despite diminished transcript levels (Supplementary Figure 1J). Collectively, these results demonstrate that close contact between T-ALL cells and tumor-associated myeloid cells is critical for leukemia survival and identify integrins as candidate mediators of these interactions.

**Figure 1.**
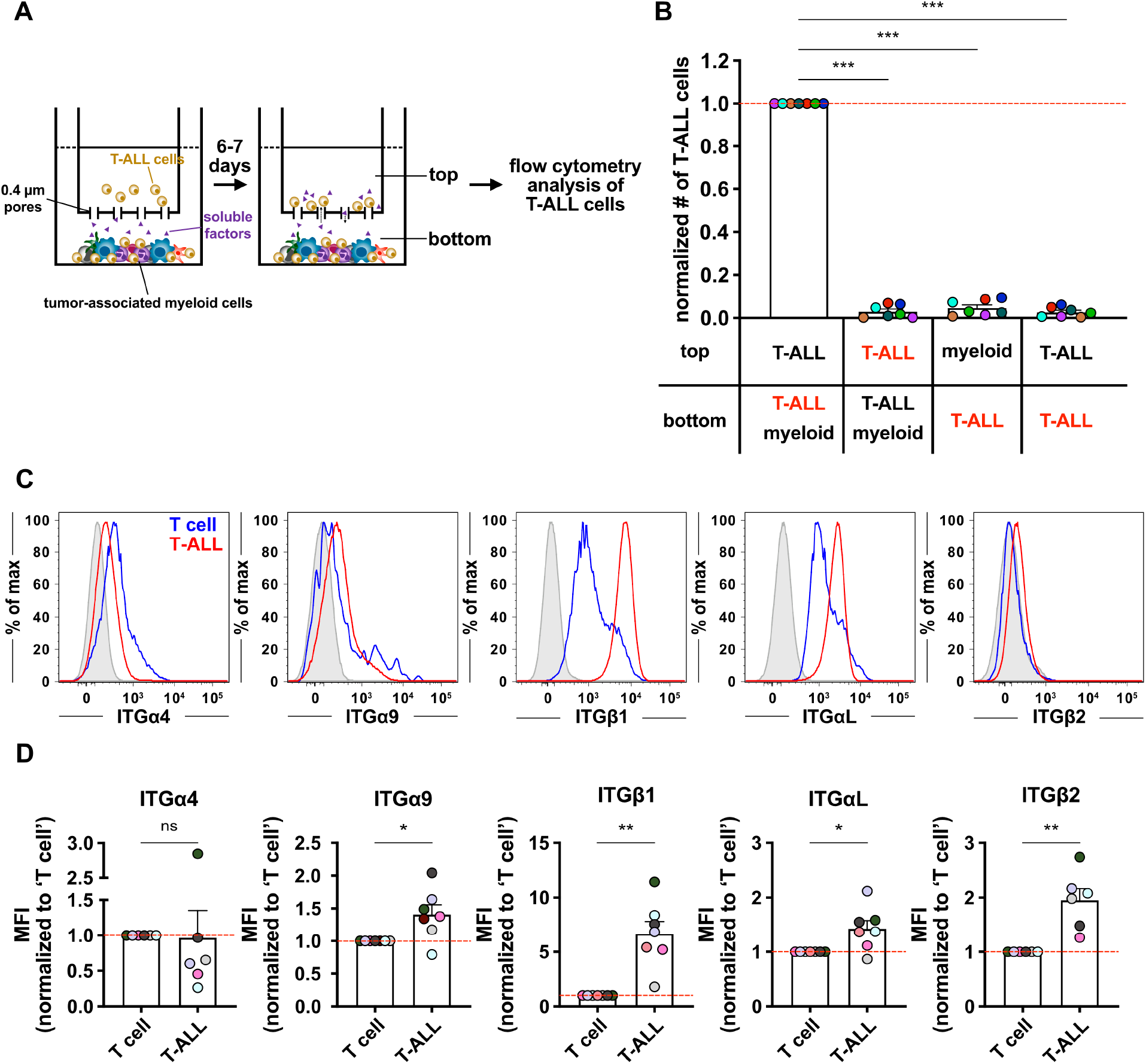
Close contact is critical for myeloid-mediated T-ALL survival, and integrin expression is elevated on T-ALL cells. (A) Schematic illustration of transwell assays in which LN3 T-ALL cells are cultured in the presence or absence of enriched tumor-associated myeloid cells in the same or opposite chambers to determine whether T-ALL cells require close contact with myeloid cells to survive. (B) Quantification of viable T-ALL cells in the chamber indicated in red 6-7 days after co-culture initiation. Results were normalized to the viability of T-ALL cells making physical contact with tumor-associated myeloid cells in the bottom chamber within each experiment. Bars depict the mean + SEM of cumulative data from 7 experiments, each with a distinct color-coded primary T-ALL; symbols represent the mean of 2-3 replicate wells per experiment. (C-D) (C) Representative flow cytometry plots and (D) cell surface integrin expression levels of transplanted LN3 T-ALL cells (red; CD45.2^+^CD5^+^) and host T cells (blue; CD45.1^+^CD5^+^) from the same leukemic spleens, quantified as median fluorescence intensity (MFI) values by flow cytometry. (C) Isotype control stains are shaded in gray. (D) Results were normalized to T-cell levels within each experiment. Data are compiled from 6-7 independent experiments, each with a distinct color-coded primary T-ALL. Symbols represent individual mice. Statistical significance was determined by (B) repeated measures one-way ANOVA with the Holm-Sidak correction, and (D) paired Student t tests; *P*-values: *<0.05, **<0.01, ***<0.001. ns, not significant.

### Inhibiting integrin-mediated cell adhesion diminishes myeloid support of T-ALL by reducing IGF1R signaling

Next, we tested whether interactions of the candidate integrins LFA-1 and/or β1 integrins with the corresponding adhesion molecules ICAM-1 and/or VCAM-1, respectively, are required for myeloid-mediated T-ALL support. First, we confirmed that ICAM-1 and VCAM-1 are expressed by tumor-associated myeloid subsets, particularly macrophages, monocytes, and cDC2, which promote T-ALL survival in vitro (16). While tumor-associated macrophages, cDC2s and monocytes all express ICAM-1, only tumor-associated macrophages express high levels of VCAM-1 (Supplementary Figure 2). LN3 T-ALL cells were then co-cultured with tumor-associated myeloid cells in the presence of anti-ICAM-1 and/or anti-VCAM-1 blocking antibodies. Blocking either ICAM-1 or VCAM-1 suppressed T-ALL survival significantly. Notably, blocking both ICAM-1 and VCAM-1 decreased survival to levels comparable to T-ALL cells cultured alone (Figure 2A, Supplementary Figure 3A-B), indicating that these adhesion molecules are essential for myeloid-mediated T-ALL support. Blocking antibodies against the corresponding integrins ITGαL and/or ITGβ1 (Figure 2B) or administration of a small molecule inhibitor to LFA-1 adhesion (Figure 2C) also significantly diminished T-ALL survival. Isotype-matched antibodies that bound T-ALL cells did not alter T-ALL viability in co-cultures with myeloid cells (Supplementary Figure 3C-D), demonstrating that blocking antibodies against integrins did not reduce T-ALL survival by inducing Fc receptor-mediated phagocytosis of opsonized T-ALL cells. These results demonstrate that tumor-associated myeloid cells support T-ALL growth through integrin-mediated close contact in vitro.

**Figure 2.**
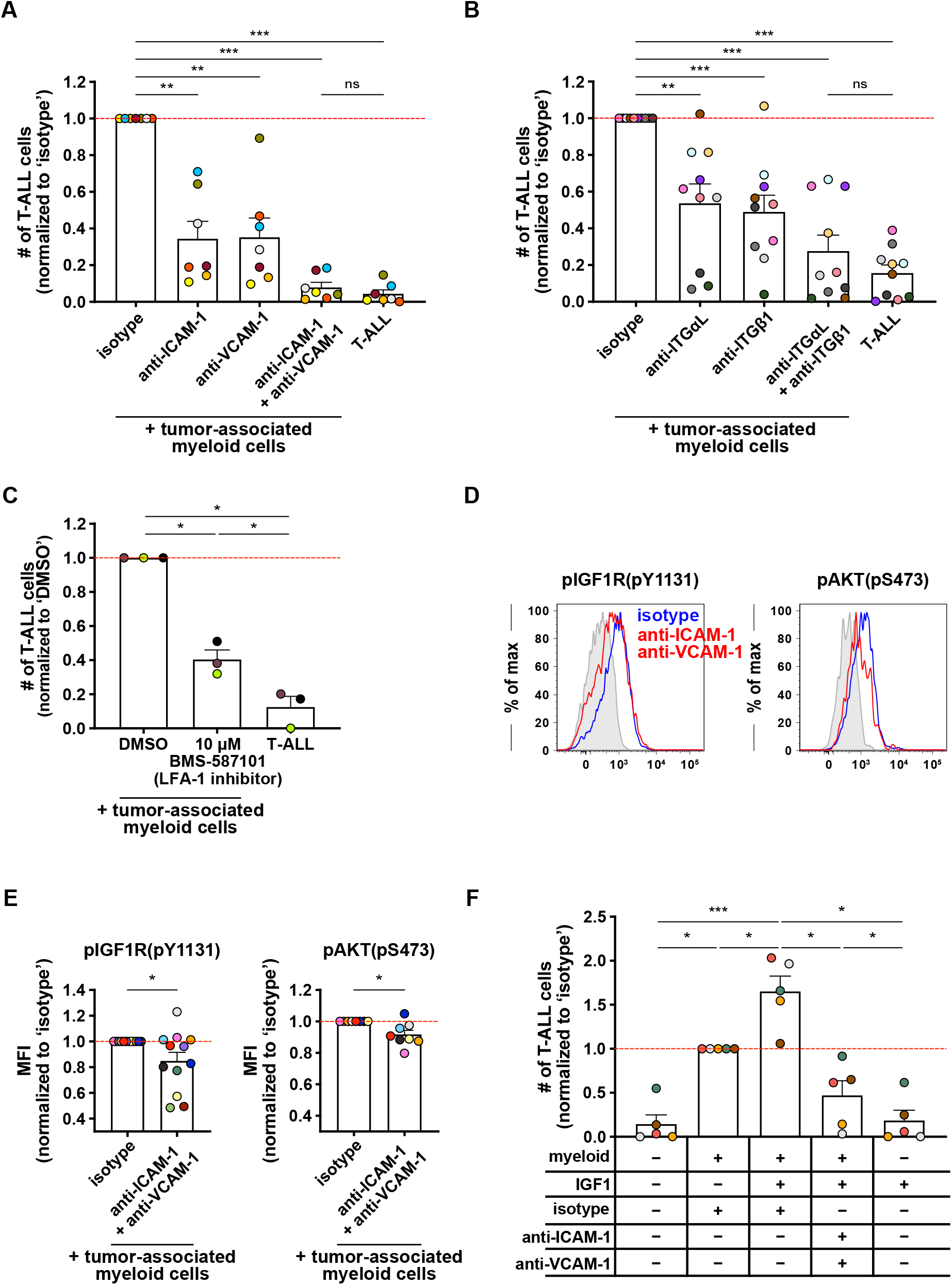
Integrin-mediated cell adhesion is required for myeloid support of T-ALL survival in vitro and for sensitization of T-ALL cells to IGF1R signaling. (A-B) Quantification of viable splenic LN3 T-ALL cells 6-7 days after co-culture with enriched tumor-associated myeloid cells in the presence of (A) anti-ICAM-1 and/or anti-VCAM-1 blocking antibodies or (B) anti-integrin αL (ITGαL) and/or anti-ITGβ1 blocking antibodies (10 µg/ml for each). Results were normalized to isotype-treated cultures. Bars show the mean + SEM from 7-10 independent experiments, each with a distinct color-coded primary T-ALL; symbols represent the average of 2-3 technical replicate wells per experiment. The red line indicates the normalized mean viability of isotype-treated T-ALL cells. (C) Quantification of viable splenic LN3 T-ALL cells 6-7 days after co-culture with enriched tumor-associated myeloid cells in the presence of 10 µM BMS-587101 (an LFA-1 inhibitor) or DMSO vehicle control. Results were normalized to DMSO-treated cultures. Bars show the mean + SEM from 3 independent experiments, each with a distinct color-coded primary T-ALL; symbols represent the average of 2-3 technical replicate wells per experiment. The red line indicates the normalized mean viability of DMSO-treated T-ALL cells. (D-E) (D) Representative flow cytometry plots of phosphorylated (p)IGF1R and the downstream signaling molecule pAKT in LN3 T-ALL cells co-cultured with tumor-associated myeloid cells for 4-5 days in the presence of anti-ICAM-1 and anti-VCAM-1 blocking antibodies (red; 10 µg/ml per each) or isotype controls (blue). Isotype control stains are shaded in gray. (E) Quantification of pIGF1R and pAKT MFI levels in co-cultured T-ALL cells, as in (D). Results were normalized to the MFI of isotype-treated cultures within each experiment. Bars represent the mean + SEM from 8-12 independent experiments, each with a distinct color-coded primary T-ALL. The red line indicates the normalized mean MFI of isotype-treated cultures. (F) Quantification of viable splenic T-ALL cells 6-7 days after co-culture with enriched tumor-associated myeloid cells in the presence or absence of exogenous IGF1 (100 ng/ml), and in the presence or absence of anti-ICAM-1 and anti-VCAM-1 blocking antibodies (10 µg/ml per each). Results were normalized to the viability of T-ALL cells co-cultured with myeloid cells in the presence of isotype control antibodies alone. Bars represent the mean + SEM of data compiled from 5 independent experiments with distinct color-coded primary T-ALL; symbols represent the average of 2-3 technical replicate wells per experiment. The red line indicates the normalized mean viability of isotype-treated co-cultures. Statistical significance was determined by (A, B, C, F) repeated measures one-way ANOVA with the Holm-Sidak correction, and (E) paired Student t tests; *P*-values: *<0.05, **<0.01, ***<0.001. ns, not significant.

**Figure 3.**
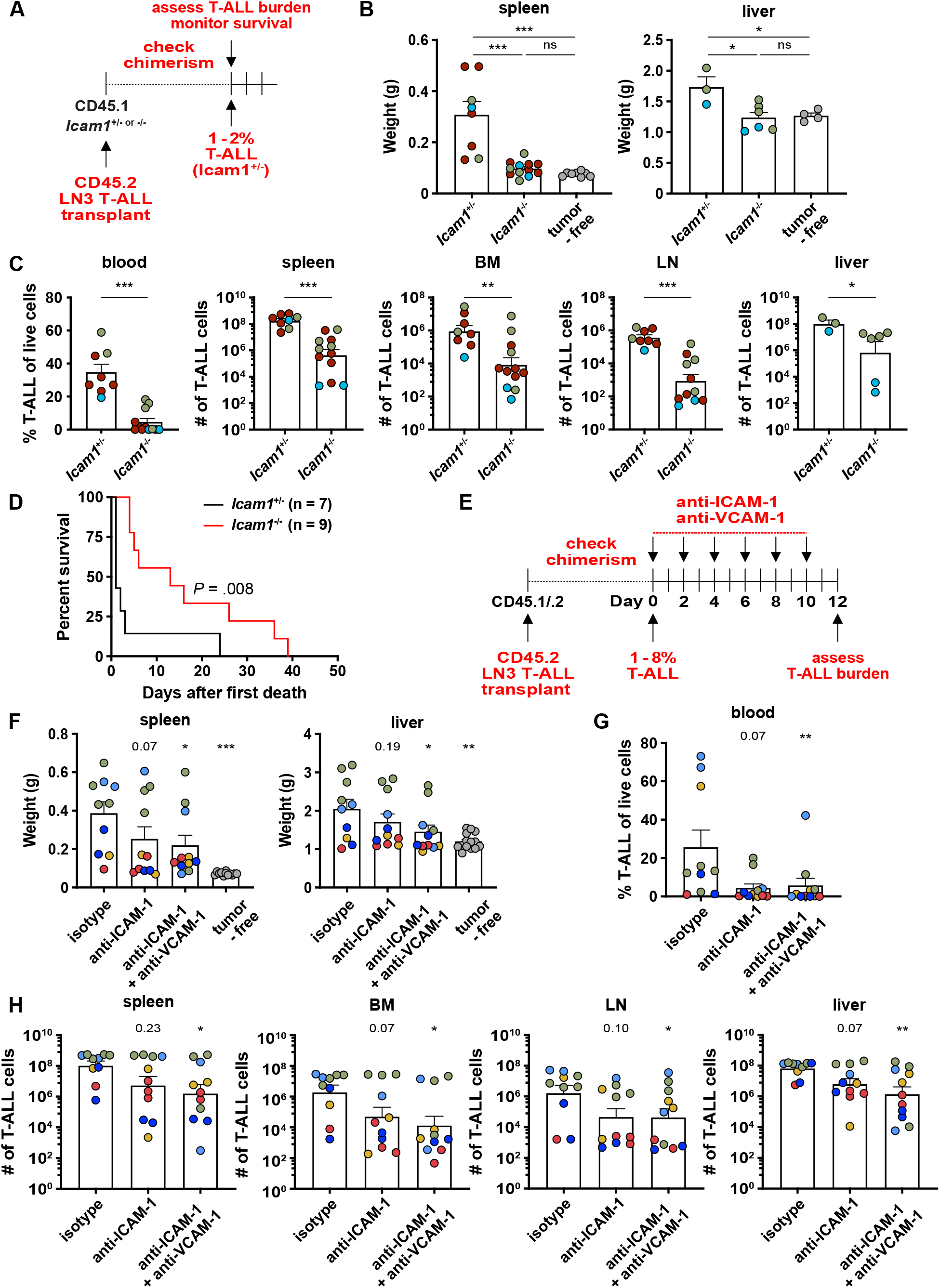
Inhibition of ICAM-1-and VCAM-1-mediated cell adhesion reduces T-ALL burden and prolongs survival in vivo. (A) Experimental schematic depicting LN3 T-ALL engraftment into *Icam1*^-/-^ and *Icam1*^+/-^ littermate controls. After T-ALL establishment (1-2% blood chimerism in controls), leukemic burden was assessed in multiple organs or survival was monitored. (B) Quantification of splenic and liver weights from *Icam1*^+/-^ and *Icam1*^-/-^ mice transplanted with primary LN3 T-ALL. Tumor-free littermate mice were included for comparison. (C) Quantification of the frequency of circulating T-ALL blasts in the blood and numbers of T-ALL cells in the spleen, bone marrow (BM), inguinal lymph nodes (LN), and liver in *Icam1*^+/-^ and *Icam1*^-/-^ mice from the same experiments as in (B). Bars depict the mean + SEM of cumulative data from 2-3 experiments, each with a distinct color-coded primary T-ALL; symbols represent individual mice. (D) Graph shows cumulative survival of *Icam1*^+/-^ and *Icam1*^-/-^ leukemic mice from 3 independent experiments, each with a different primary LN3 T-ALL, using the strategy as in (A). The Kaplan-Meier survival curves were normalized to the first day of death in each experiment. (E) Experimental schematic depicting dosing schedule for anti-ICAM-1, anti-VCAM-1, or isotype control antibodies following establishment of LN3 T-ALL (>1% spleen chimerism) in congenic hosts. Leukemia burden was assessed two days after the final antibody injection. (F) Quantification of spleen and liver weights from the experiments depicted in (E). Tumor-free, age-matched mice were included for comparison. Bars depict the mean + SEM of cumulative data from 5 experiments, each with a distinct color-coded primary T-ALL; symbols represent individual mice. (G-H) Quantification of the (G) frequency of circulating T-ALL blasts in the blood and (H) number of T-ALL cells in the spleen, BM, LN, and liver from the same experiments as in (F). Statistical significance was determined by (B, F, G, H) repeated measures one-way ANOVA with the Holm-Sidak correction, (C) unpaired Student t tests, and (D) log-rank tests; *P*-values: *<0.05, **<0.01, ***<0.001. ns, not significant.

We previously found that tumor-associated myeloid cells support T-ALL growth by activating IGF1R signaling (16), and crosstalk has been shown between integrin and IGF1R signaling (25,29,30). Thus, we hypothesized that inhibiting integrin adhesion would diminish myeloid-mediated IGF1R activation in T-ALL cells. Indeed, blockade of ICAM-1 and VCAM-1 significantly diminished the levels of activated, phosphorylated (p)IGF1R and the downstream kinase pAKT (Figure 2D-E). Addition of exogenous IGF1 enhances survival of T-ALL cells co-cultured with tumor-associated myeloid cells (16). To determine whether blockade of integrin-mediated physical interactions would prevent sensitization of T-ALL cells to IGF1, we co-cultured T-ALL cells with myeloid cells in the presence or absence of exogenous IGF1 and/or antibodies against adhesion molecules. Consistent with our previous findings, addition of IGF1 increased viability of T-ALL cells only in the presence of tumor-associated myeloid cells. Notably, blockade of ICAM-1 and VCAM-1 significantly reduced sensitization of T-ALL cells to IGF1 (Figure 2F), demonstrating that integrin activation is required for the full responsiveness of T-ALL cells to IGF1. Taken together, integrin-mediated close contact between T-ALL cells and tumor-associated myeloid cells plays an essential role in promoting the survival of T-ALL by enabling activation of IGF1R signaling.

### ICAM-1-and VCAM-1-mediated cell adhesion is critical for T-ALL progression in vivo

To determine whether integrin-mediated adhesion promotes T-ALL establishment or progression in vivo, we engrafted primary LN3 T-ALL cells into *Icam1*^-/-^ mice and *Icam1*^+/-^ littermate controls (Supplementary Figure 4A) and assessed leukemic burden after T-ALL establishment in control mice (1-2% blood chimerism; Figure 3A). In *Icam1*^*-/-*^ mice, spleen and liver weights were reduced to levels comparable to tumor-free littermate *Icam1*^*+/-*^ controls (Figure 3B). Moreover, circulating T-ALL blasts were significantly reduced, as was T-ALL burden in multiple organs including the spleen, bone marrow (BM), inguinal lymph nodes (LN), and liver (Figure 3C, Supplementary Figure 4B). In keeping with these findings, *Icam1* deficiency prolonged survival following transplantation of T-ALL (Figure 3D). In order to have a sufficient number of littermates for these experiments, we compared T-ALL burden and survival in *Icam1*^*-/-*^ versus *Icam1*^*+/-*^ mice. We found that haploinsufficiency of *Icam1* resulted in slightly diminished expression of ICAM-1 on myeloid cells relative to wild-type *Icam1*^*+/+*^ cells (Supplementary Figure 4A). Thus, the reduced T-ALL burden and increased survival in *Icam1*^*-/-*^ *versus Icam1*^*+/-*^ mice could underestimate the role of ICAM-1 in supporting T-ALL survival and progression. Together, these results indicate that ICAM-1-mediated integrin activation is critical for T-ALL establishment and/or progression.

**Figure 4.**
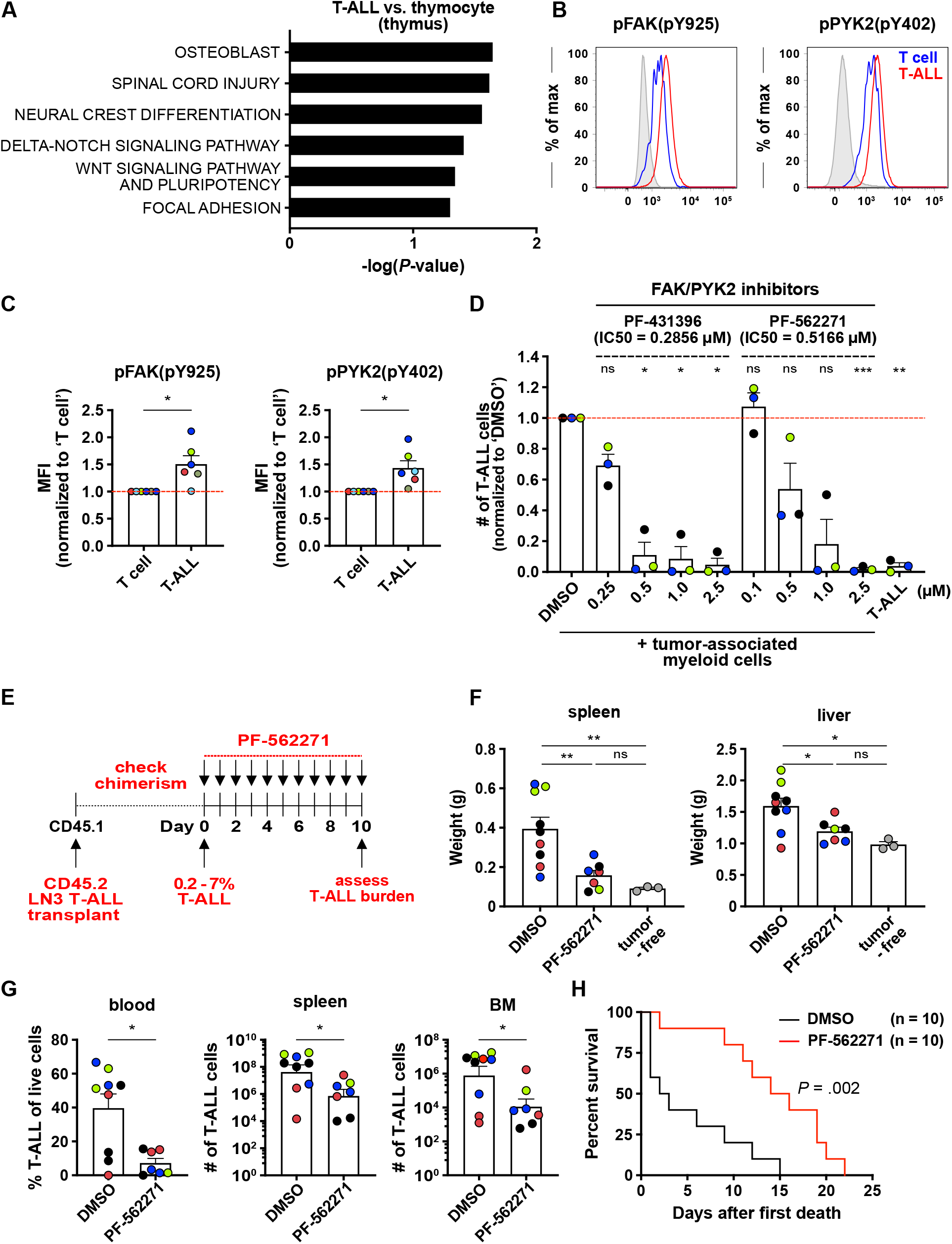
Inhibition of FAK and PYK2 signaling reduces T-ALL survival in vitro, diminishes leukemic burden in vivo, and prolongs survival. (A) The top 6 pathways preferentially activated in thymic T-ALL cells relative to healthy thymocytes were identified using the WikiPathways 2019 Mouse gene sets in the Enrichr bioinformatics tool. (B-C) (B) Representative flow cytometry plots showing levels of Pfak and pPYK2, two key kinases downstream of integrin signaling, in transplanted LN3 T-ALL cells (red) and host T cells (blue) from the same leukemic spleens. Isotype control stains are shaded in gray. (C) Quantification of the MFIs of pFAK and pPYK2 in T-ALL cells (CD45.2^+^CD5^+^) relative to host T cells (CD45.1^+^CD5^+^), as in (B). Results were normalized to MFIs of T cells within each experiment. Bars represent the mean + SEM from 6 independent experiments, each with a distinct color-coded primary T-ALL. The red line indicates the normalized mean MFI in T cells. (D) Quantification of viable T-ALL cells co-cultured with enriched tumor-associated myeloid cells for 4 days before addition of the indicated concentrations of FAK/PYK2 dual inhibitors (PF-431396 or PF-562271) or vehicle control (DMSO). The number of viable T-ALL cells was assessed after 2-3 days. Results were normalized to DMSO vehicle-treated cultures in each experiment. Bars represent means + SEM from 3 independent experiments using distinct color-coded primary T-ALLs; symbols represent the average of 2-3 technical replicate wells. The red line indicates the normalized mean T-ALL viability in DMSO-treated cultures. (E) Experimental schematic depicting dosing schedule for PF-562271 or DMSO vehicle control treatment to inhibit FAK and PYK2 signaling in mice following establishment of LN3 T-ALL in congenic recipients (>0.2% spleen chimerism). (F) Quantification of spleen and liver weights from mice transplanted with primary LN3 T-ALL after treatment with PF-562271 or DMSO, as in (E). Tumor-free littermates were included for comparison. (G) Quantification of the frequency of circulating T-ALL blasts in the blood and the number of T-ALL cells in the spleen and BM from the same experiments as in (F). Bars depict the mean + SEM of cumulative data from 4 experiments, each with a distinct color-coded primary T-ALL; symbols represent individual mice. (H) Graph shows cumulative survival of PF-562271 or DMSO-treated leukemic mice from 3 independent experiments, each with a different primary LN3 T-ALL, using the strategy as in (E). Kaplan-Meier survival curves were normalized to the first day of death in each experiment. Statistical significance was determined by (A) Fisher exact tests, (C, G) unpaired Student t tests, (D, F) repeated measures one-way ANOVA with the Holm-Sidak correction, and (H) log-rank tests; *P*-values: *<0.05, **<0.01, ***<0.001. ns, not significant.

To evaluate the contribution of integrin-mediated cell adhesion in T-ALL progression after tumor establishment, we administered an anti-ICAM-1 blocking antibody alone or in combination with an anti-VCAM-1 blocking antibody after LN3 leukemic burden reached 1-8% in the spleen (Figure 3E). Consistent with in vitro results, combination treatment using anti-ICAM-1 and anti-VCAM-1 antibodies significantly diminished circulating T-ALL blasts and T-ALL burden by one to two orders of magnitude in the spleen, BM, LN, and liver, with a similar trend observed for mice treated with anti-ICAM-1 antibody alone (Figure 3F-H, Supplementary Figure 4C). Notably, these antibodies did not deplete myeloid cells in leukemic mice (Supplementary Figure 4D-F), indicating that the observed reduction of T-ALL burden resulted from inhibition of cell adhesion, rather than from depletion of leukemia-supportive myeloid cells. Collectively, ICAM-1- and VCAM-1-mediated cell adhesion play a critical role in promoting T-ALL growth and progression in vivo.

### Myeloid cells activate FAK/PYK2 signaling to promote T-ALL survival and progression

Blocking multiple integrins and/or adhesion molecules simultaneously could be therapeutically challenging (43,44); thus, we sought a common targetable signal downstream of integrin activation. Our transcriptional profiling data from thymic T-ALL vs healthy thymocytes revealed that the “focal adhesion” pathway was among the top pathways preferentially activated in T-ALL cells (*P* = 0.049; Figure 4A). FAK and PYK2 are two integrin-associated kinases in the focal adhesion pathway that have been shown to play important roles in tumor progression (45). Thus, we hypothesized that these kinases would be activated by myeloid-mediated integrin signaling in T-ALL cells and would contribute to leukemia growth. Indeed, levels of pFAK and pPYK2 were significantly elevated in T-ALL cells relative to T cells from leukemic spleens (Figure 4B-C). Furthermore, addition of blocking antibodies to ICAM-1 and VCAM-1 in co-cultures of T-ALL and myeloid cells diminished the frequency of T-ALL cells with activated FAK or PYK2 (Supplementary Figure 5A-B), confirming a link between integrin signaling and FAK/PYK2 activation in T-ALL cells upon contact with myeloid cells. We next tested whether activation of FAK and PYK2 is required for myeloid-mediated support of T-ALL. In co-cultures, small molecule inhibitors of FAK/PYK2 significantly reduced LN3 T-ALL survival in a concentration dependent manner (Figure 4D). Notably, administration of a FAK/PYK2 inhibitor (PF-562271) in leukemic mice (Figure 4E) diminished spleen and liver weights to those of healthy littermate controls (Figure 4F). FAK/PYK2 inhibitor treatment also significantly reduced circulating T-ALL blasts and tumor burden by one to two orders of magnitude in the spleen and BM (Figure 4F-G, Supplementary Figure 5C). Furthermore, in vivo administration of this FAK/PYK2 inhibitor significantly prolonged survival of mice with established LN3 T-ALL (Figure 4H). PF-562271 altered not only the T-ALL cells, but also the TME, consistent with previous findings (46). The number of cDC1, cDC2, and Gr1^lo^ monocytes was reduced, but the overall number of myeloid cells and the number of macrophages did not significantly decline (Supplementary Figure 5D-F). Given that macrophages in the splenic and hepatic TME have the most potent capacity among myeloid subsets to support T-ALL (16), the observed reduction of leukemic burden and prolonged survival likely result mainly from inhibiting FAK/PYK2 signaling in T-ALL cells, not from altered myeloid composition; however, the ability of these inhibitors to diminish leukemia-supportive myeloid cells could be an added benefit of targeting this pathway.

**Figure 5.**
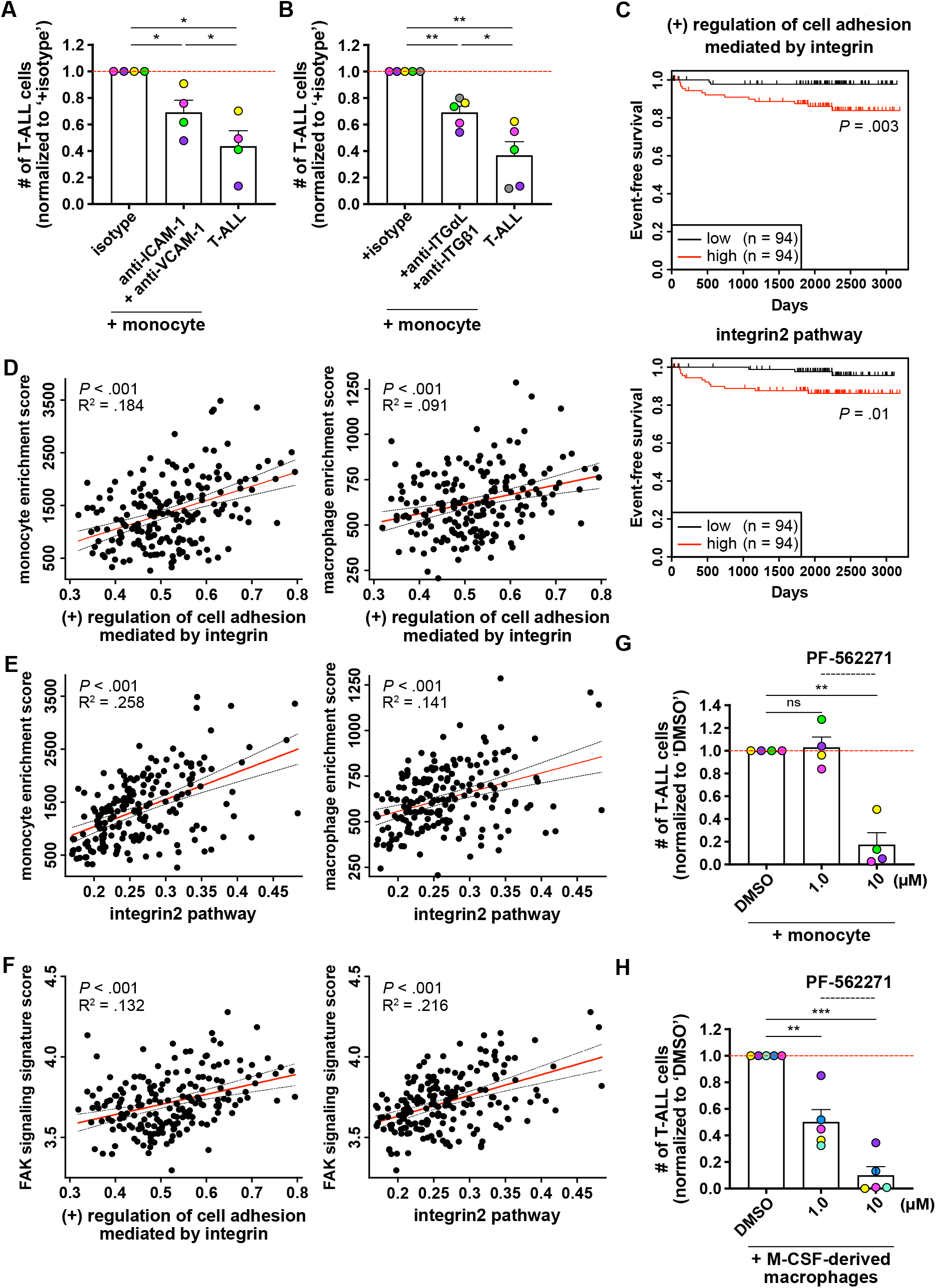
Human myeloid cells promote survival of primary patient T-ALL cells in an integrin-dependent manner in vitro, and enriched gene signatures of integrin pathways in patients are associated with worse prognosis. (A-B) Quantification of viable primary patient T-ALL cells cultured for 6-7 days alone or with monocytes from healthy donor peripheral blood mononuclear cells (PBMCs). Co-cultures were carried out in the presence of (A) anti-ICAM-1 (20 µg/ml) and anti-VCAM-1 (10 µg/ml) blocking antibodies, (B) anti-integrin αL (ITGαL) and anti-ITGβ1 blocking antibodies (10 µg/ml for each), or isotype control antibodies as indicated. Results were normalized to isotype-treated cultures in each experiment. Bars represent means + SEM from 2-3 independent experiments using 4-5 distinct, color-coded patient-derived T-ALLs; symbols represent the average of 2 technical replicate wells. The red line indicates the normalized mean T-ALL viability in isotype-treated cultures. (C) Longitudinal event-free survival is plotted for pediatric T-ALL patients stratified into two equal groups based on their gene signature enrichment scores for “(+) regulation of cell adhesion mediated by integrin” (top) or “integrin2 pathway” (bottom). Patient data were analyzed from published datasets from 264 T-ALL patients from the TARGET ALL Phase II trial (14). (D-E) Plots depict correlations between enrichment scores of the gene signatures (D) “(+) regulation of cell adhesion mediated by integrin” or (E) “integrin2 pathway” with enrichment scores for monocytes (left) and macrophages (right) in patient samples. The red and black dotted lines represent the best-fit line and 95% confidence bands, respectively. Symbols represent each patient. (F) Plots show the correlation between enrichment scores for the gene signatures “(+) regulation of cell adhesion mediated by integrin” (left) or “integrin2 pathway” (right) with signature scores of FAK signaling in patient samples. The enrichment scores for “FAK pathway” were calculated using the gene signatures from a prior study (66). The red and black dotted lines represent the best-fit line and 95% confidence bands, respectively. Symbols represent each patient. (G-H) Quantification of viable primary patient T-ALL cells co-cultured for 4 days with (G) PBMC-derived monocytes or (H) M-CSF-derived macrophages 2-3 days after addition of the indicated concentrations of PF-562271 or vehicle control (DMSO). Results were normalized to DMSO-treated cultures for each experiment. Bars represent means + SEM from 3 independent experiments using 4-5 distinct, color-coded patient-derived T-ALLs; symbols represent the average of 2 technical replicate wells. The red line indicates the normalized mean T-ALL viability in DMSO-treated cultures. Statistical significance was determined by (A, B, G, H) repeated measures one-way ANOVA with the Holm-Sidak correction, (C) log-rank tests, and (D, E, F) simple linear regression analyses; *P*-values: *<0.05, **<0.01, ***<0.001. ns, not significant.

To determine whether integrin-mediated myeloid support of T-ALL was relevant in a distinct T-ALL mouse model, we carried out comparable experiments in human CD2 promoter-driven LIM domain only 2 (LMO2) transgenic mice (47). As in LN3 T-ALL, LMO2 T-ALL cells express higher levels of Itgα9 and Itgβ1 transcripts and proteins and higher levels of LFA-1 protein relative to non-leukemic T cells from the same splenic tumor environment (Supplementary Figure 6A-B). In addition, integrin-mediated cell adhesion promotes survival of LMO2 T-ALL cells co-cultured with tumor-associated myeloid cells (Supplementary Figure 6C). Moreover, inhibition of FAK/PYK2 significantly reduces survival of T-ALL cells co-cultured with tumor-associated myeloid cells (Supplementary Figure 6D). Taken together, these concordant findings in multiple mouse models of T-ALL demonstrate that tumor-associated myeloid cells promote T-ALL survival and growth via activation of integrin signaling and downstream FAK/PYK2 kinases.

### Integrin activation by myeloid cells promotes primary patient T-ALL survival and is associated with inferior outcomes

We previously demonstrated that human peripheral blood mononuclear cell (PBMC)-derived myeloid cells support survival of primary patient T-ALL cells (16), and thus hypothesized a role for integrin-mediated cell adhesion in human T-ALL. We first assessed expression of integrins and cell adhesion ligands by T-ALL cells and PBMC-derived myeloid cells, respectively. Consistent with mouse models, VLA-4, integrin α9β1, and LFA-1 were expressed by patient T-ALL cells (Supplementary Figure 7A). ICAM-1 was highly expressed by both monocytes and M-CSF-derived macrophages, but VCAM-1 expression was negligible (Supplementary Figure 7B-C). To determine the impact of integrin-mediated cell adhesion on myeloid support of T-ALL, primary pediatric T-ALL cells were co-cultured with monocytes or M-CSF-derived macrophages in the presence or absence of blocking antibodies against ICAM-1 and VCAM-1. Blocking adhesion molecules or integrins significantly diminished survival of T-ALL cells co-cultured with monocytes (Figure 5A-B), but not with M-CSF-derived macrophages (Supplementary Figure 7D). Isotype-matched antibodies that bound T-ALL cells did not reduce T-ALL survival in myeloid co-cultures (Supplementary Figure 7E-F), indicating that Fc receptor-mediated phagocytosis of opsonized T-ALL cells was not a significant factor in reduced T-ALL viability after treatment of T-ALL cells with anti-integrin antibodies. To test whether integrin signaling is associated with altered patient outcomes, we analyzed published datasets from pediatric T-ALL patients (14) and found that increased gene signatures of the “(+) regulation of cell adhesion mediated by integrin” and “integrin2” pathways correlate significantly with diminished event-free survival (Figure 5C). Also, elevated expression of integrin pathway gene signatures correlates significantly with monocyte and macrophage enrichment scores in T-ALL patient samples (Figure 5D-E), suggesting myeloid cells activate integrin signaling in patient T-ALL cells. Furthermore, integrin and FAK pathway signature scores significantly correlate with one another (Figure 5F), concordant with a link between integrin and FAK signaling in T-ALL patients. Notably, survival of T-ALL cells co-cultured with monocytes or M-CSF-derived macrophages was significantly impaired in the presence of a FAK/PYK2 inhibitor (PF-562271; Figure 5G-H), implicating a critical role for integrin activation and downstream FAK/PYK2 signaling in T-ALL progression. Collectively, these results indicate that myeloid cells support survival of patient T-ALL cells via integrin-mediated cell adhesion and downstream activation of FAK/PYK2.

## Discussion

Although tumor-associated myeloid cells promote T-ALL progression by activating IGF1R signaling (15,16), which is required for initiation and growth of T-ALL (48), exogenous IGF1 alone is not sufficient to support T-ALL survival (16). Notably, the presence of tumor-associated myeloid cells enhances the response of T-ALL cells to exogenous IGF1 (16). These findings suggest that myeloid cells employ mechanisms in addition to IGF1 production to promote T-ALL survival in the TME. Analysis of our published RNA-seq datasets (16) revealed that gene expression signatures of cell-cell interactions were significantly enriched in thymic T-ALL cells relative to healthy thymocytes, consistent with our previous imaging results showing that T-ALL cells make prolonged and frequent contacts with DCs in the thymus (15). Based on these findings, we hypothesized that physical interactions are critical for myeloid-mediated T-ALL support. Here we find that T-ALL cells express elevated levels of integrins known to be involved in T cell-myeloid interactions (24). Moreover, inhibition of integrin-mediated cell adhesion results in a significant reduction in T-ALL cellularity both in vitro and in vivo and prolongs survival in a mouse model of T-ALL. In other cancers, integrins have been shown to interact with IGF1R to promote signaling that results in tumor survival, migration, and therapeutic resistance (30,49–51). Notably, we found that blocking integrin-mediated cell adhesion diminishes levels of activated IGF1R and AKT in T-ALL cells co-cultured with tumor-associated myeloid cells, and also impairs myeloid-mediated sensitization of T-ALL cells to exogenous IGF1. These findings link the dual requirements of integrin activation and IGF1R signaling for myeloid-mediated support of T-ALL survival. Consistent with mouse models, survival of primary patient T-ALL cells co-cultured with monocytes was significantly reduced by blocking integrin-mediated cell adhesion. Also, pediatric T-ALL patients with elevated expression of integrin pathway genes have increased gene signatures associated with myeloid cells and inferior outcomes. Altogether, these results indicate that myeloid cells promote T-ALL survival through a combination of close contact with leukemia cells, driven by integrin-mediated interactions and IGF1R activation.

In keeping with a critical role for integrins in T-ALL survival, FAK and PYK2 signaling contributes to myeloid-mediated T-ALL support. FAK and PYK2 promote tumor initiation, growth, and invasion across multiple malignancies (37,45,52), and FAK has been reported to promote survival of *PTEN*-null T-ALL cells (53). We found that FAK and PYK2 are highly activated in T-ALL cells relative to T cells from the same leukemic environment, consistent with elevated integrin signaling in T-ALL. Given that FAK and PYK2 are activated downstream of integrin signaling and promote T-cell proliferation and survival (26), we hypothesized that myeloid-mediated T-ALL support would be dependent on FAK and/or PYK2 signaling. Indeed, pharmacologic inhibition of these kinases impairs myeloid-mediated T-ALL support in a dose-dependent manner in vitro, reduces T-ALL burden in vivo, and prolongs survival of leukemic mice, pointing to FAK and PYK2 as potential therapeutic targets for T-ALL.

FAK and PYK2 can be activated not only by integrins, but also by soluble factors, including vascular endothelial growth factor (VEGF), the chemokine CCL19, and interleukin-1β (IL-1b) (54–56). When mouse or patient T-ALL cells were co-cultured with tumor-associated or PBMC-derived myeloid cells, respectively, they exhibited fairly uniform sensitivity to FAK/PYK2 dual inhibitors, whereas the efficacy of inhibiting integrin-mediated cell adhesion varied between distinct primary T-ALLs. These findings suggest that tumor-associated myeloid cells may activate FAK and PYK2 in T-ALL cells not only through integrin signaling, but also through additional pathways. A previous study reported that *Vegfa* and *Il1b* transcripts are elevated in leukemia-associated macrophages in a NOTCH1-induced model of T-ALL (57), suggesting secreted myeloid factors could promote FAK/PYK2 activation in T-ALL cells. Thus, a better understanding of the mechanisms by which tumor-associated myeloid cells activate FAK and PYK2 signaling and how these pathways contribute to T-ALL progression could reveal therapeutic opportunities in T-ALL.

Although inhibiting integrin adhesion or FAK/PYK2 signaling significantly diminished leukemia burden and prolonged survival in a mouse model of T-ALL, the mice eventually succumbed to leukemia, suggesting other cell-intrinsic pathways or supportive elements of the TME would need to be inhibited simultaneously for therapeutic intervention. Previous studies have identified some other signals in the TME that support T-ALL. For example, despite activating mutations in *NOTCH1*, T-ALL cells require NOTCH ligands, which are expressed by non-hematopoietic stromal cells in a variety of organs (27,58), to activate NOTCH1 (11,48,59). IL-7, which is produced by TECs, has also been implicated in promoting T-ALL progression (21,22). IGF1 is produced in the liver as a growth factor but interestingly, we have found it is also expressed by leukemia-associated myeloid cells, and IGF1R signaling is critical for T-ALL initiation and growth (15,16,48). Furthermore, CXCL12 expression by BM stromal cells has been shown to promote T-ALL initiation and progression (19,20). Cell adhesion molecules such as integrins and CD44 have also been implicated in T-ALL pathogenesis through interactions with BM stromal cells (38,39,60). Thus, as our study reveals a critical role for integrins in myeloid-mediated support of T-ALL, it will likely be important to target signals provided by hematopoietic and other stromal cells in the TME to improve therapeutic strategies for T-ALL.

Collectively, this study demonstrates that tumor-associated myeloid cells support T-ALL through integrin-mediated close contact. Inhibition of integrins or downstream FAK/PYK2 signaling delays T-ALL progression and improves survival. Integrins are associated with T-cell localization to and migration within secondary lymphoid organs, such as LNs where pro-leukemic cytokines like IL-7 are produced (22,61). Moreover, integrin-mediated migration and adhesion is activated by chemokine receptors like CXCR4, which promotes T-ALL initiation and maintenance in the BM (19,20,62). Thus, inhibition of integrin-mediated physical interactions could target multiple mechanisms by which the TME supports T-ALL. It will be important to evaluate whether targeting integrins or downstream FAK and PYK2 signaling along with chemokine or growth factor receptor pathways will prove therapeutically efficacious alone or in the context of conventional chemotherapeutic approaches.

## Methods

### Mice

LN3 (gift from T. Serwold) (40), CD2-*Lmo2* transgenic (LMO2; gift from U. Davé) (47), B6.129S4-*Icam1*^*tm1Jcgr*^/J (*Icam1*^-/-^) (63), C57BL/6J, and B6.SJL-*Ptprc*^*a*^ *Pepc*^*b*^/BoyJ (CD45.1) mouse strains were bred in-house and housed in specific-pathogen-free conditions. Mice were sourced from The Jackson Laboratory, unless otherwise noted. All experimental procedures were approved by the Institutional Animal Care and Use Committee at the University of Texas at Austin.

### T-ALL transplant model

For all cellular assays, a total of 5×10^6^ primary T-ALL cells, isolated from spleens of leukemic CD45.2^+^ LN3 or LMO2 mice, were resuspended in sterile phosphate-buffered saline (PBS), and injected intraperitoneally (i.p.) into nonirradiated sex-matched CD45.1 *Icam1*^-/-^, CD45.1, or CD45.1/CD45.2 mice, as indicated. T-ALL engraftment (CD45.2^+^CD5^+^) in the spleens or tail blood was evaluated by flow cytometry to confirm tumor establishment.

### Flow cytometric analysis

See Supplementary Methods for experimental details.

### Transwell assays

See Supplementary Methods for experimental details.

### In vitro cultures of mouse and human T-ALL cells

See Supplementary Methods for experimental details.

### In vivo inhibition of integrin-mediated cell adhesion or FAK/PYK2 signaling

*Icam1*^-/-^ or *Icam1*^+/-^ littermate controls (6-8 weeks old) were engrafted i.p. with 5×10^6^ primary LN3 T-ALL cells. At 1-2% blood chimerism in *Icam1*^+/-^ mice, T-ALL burden was assessed. Survival was monitored in separate cohorts.

For antibody-mediated blockade of adhesion molecules, congenic mice with established T-ALL burden (>1% splenic chimerism) were randomly divided into three groups and treated i.p. with 100 µg each of *InVivo*MAb antibodies in 200 µl of PBS every 2 days for a total of 5 injections: 1) anti-mouse CD54 (YN1/1.7.4) alone, 2) a cocktail of anti-mouse CD54 and CD106 (M/K-2.7), or 3) respective isotype controls (HRPN and LTF-2; all InVivoMAb antibodies are from BioXCell). T-ALL burden was evaluated 24-48 hours after the final injection.

For inhibition of FAK/PYK2 signaling, congenic mice with established T-ALL burden (>0.2% splenic chimerism) were randomly divided into treated and control groups. PF-562271 (Selleckchem) (64,65) was administered i.p. at 25 mg/kg body weight in 200 ul injection solution once daily for 10-11 consecutive days. Injection solution was comprised of 10% PF-562271 (or DMSO), 40% PEG300 (Selleckchem), 5% TWEEN 80 (Sigma-Aldrich), and 45% PBS. T-ALL burden was evaluated 1-3 hours after the final injection. All comparisons within experiments were carried out with age-matched mice (6-8 weeks old) engrafted with the same primary LN3 T-ALL.

### Quantitative PCR

See Supplementary Methods for experimental details.

### Bioinformatic analysis of mouse and patient T-ALL RNA-seq data

See Supplementary Methods for experimental details.

### Statistical analysis

Statistical analyses were performed with Prism (GraphPad Software). Normality was determined using the D’Agostino & Pearson or Shapiro-Wilk tests, as appropriate for sample size. Statistical significance was determined using unpaired Student t tests for normally distributed data or the non-parametric Mann-Whitney test, paired Student t tests, repeated measures one-way ANOVA with the Holm-Sidak correction for normally distributed data or the non-parametric Friedman test, two-way ANOVA with the Holm-Sidak correction, or Log-rank tests, as indicated in figure legends.

## Data availability

The data generated in this study are available within the article and its supplementary data files. The mouse RNA-seq data analyzed in this study were obtained from Gene Expression Omnibus (GEO) at GSE150096. The patient RNA-seq data analyzed in this study were obtained from the TARGET website at https://ocg.cancer.gov/programs/target. For original data, please contact.

## Acknowledgements

We thank all members of the Ehrlich lab for helpful discussion and advice. We also thank the staff of the animal facility and the Center for Biomedical Research Support at the University of Texas at Austin for technical assistance. This work was supported by RP180073 from the Cancer Prevention and Research Institute of Texas (to L.I.R.E.), RSG367515 from the American Cancer Society (to L.I.R.E.), and the William and Ella Owens Medical Research Foundation (to L.I.R.E.).

## Authors’ Contributions

A.L., S.H.N., and R.S.H. performed experiments; T.A.D., Z.H., and D.A. carried out data analyses; A.L. designed experiments, analyzed results, generated figures, and wrote the manuscript; T.M.H. provided advice and materials; L.I.R.E. designed experiments, analyzed results, and wrote the manuscript.

