## Supplementary materials for "Integrin signaling is critical for myeloid-mediated support of T-cell acute lymphoblastic leukemia"

#### Supplementary Methods

##### Flow cytometric analysis

Organs were harvested and processed as follows: Spleens were mechanically dissociated with FACS wash buffer (FWB: phosphate-buffered saline (PBS) supplemented with 2% (v/v) BCS (Bovine Calf Serum; GemCell) and 0.5 mM EDTA). Bone marrow cells were obtained by flushing femurs with 2 ml of FWB. Inguinal lymph nodes were enzymatically digested with a cocktail of 0.6 mg/ml (w/v) Liberase and 20 U/ml DNase I (both from Roche). LNs were sequentially digested 3 times with 2 ml of cocktail for 12 minutes per digest. Livers were mechanically dissociated, spun at 50 g for 3 minutes, and the supernatant was collected and analyzed. All single-cell suspensions were filtered using 40 µm filters (Fisher Scientific) and subjected to red blood cell lysis using RBC Lysis Buffer (BioLegend). Cells were immunostained by incubating at 4˚C for 30 minutes with fluorescently labeled antibodies below (all antibodies were purchased from BioLegend, unless otherwise indicated). After staining, cells were washed 1-2 times in FWB and resuspended in FWB containing 1 µg/ml propidium iodide (PI; Enzo Life Sciences) to assess viability. All flow cytometric data were acquired using an LSR II flow cytometer (BD Biosciences). Post-acquisition data analysis was performed using FlowJo (v9.9.6, Tree Star, Inc.).

For mouse lymphoid staining, anti-mouse CD4-PerCP/Cy5.5 (RM4-5), CD5-PE (53-7.3), CD8-Pacific Blue (53-6.7), CD45.1-FITC (A20), CD45.2-APC/Cy7 (104), B220-Alexa Fluor 700 (RA3-6B2), and TCRβ-PE/Cy7 (H57-597) antibodies were used. For mouse myeloid staining, CD64-PE (X54-5/7.1), CD11c-Pacific Blue (N418), I-A/I-E-APC/Cy7 (M5/114.15.2), CD172a-PE/Cy7 (SIRPα; P84), biotinylated XCR1 (ZET), CD11b-Alexa Fluor 700 (M1/70), CD115-Alexa Fluor 488 (AFS98), Gr1-Brilliant Violet 510 (RB6-8C5), F4/80-Alexa Fluor 647 (T45-2342; BD Biosciences), and Qdot 605 streptavidin conjugate (Invitrogen) antibodies were used. For mouse integrin staining, anti-mouse CD11a-FITC (ITGαL; 2D7), CD18-PE (ITGβ2; M18/2), CD29-PE (ITGβ1; HMβ1-1), CD49d-Alexa Fluor 488 (ITGα4; R1-2), Integrin α9 (N-19; Santa Cruz Biotechnology), and donkey anti-goat IgG-Alexa Fluor 488 (Jackson ImmunoResearch) antibodies were used. For mouse adhesion molecule staining, anti-mouse CD54-PerCP/Cy5.5 (ICAM-1; YN1/1.7.4) and CD106-PerCP/Cy5.5 (VCAM-1; 429(MVCAM.A)) antibodies were used. The relevant isotype antibodies (RTK2758, RTK4530, HTK888, and Poly24030) were used as controls.

For human myeloid staining, anti-human CD11b-Alexa Fluor 700 (M1/70) and CD14-PerCP/Cyanine5.5 (M5E2) were used. For human integrin staining, anti-human CD11a-PE (HI111), CD18-FITC (TS1/18), CD29-Alexa Fluor 488 (TS2/16), CD49d-PE (9F10), and Integrin α9β1-PE (Y9A2) antibodies were used. For human adhesion molecule staining, anti-human CD54-FITC (HA58), CD106-APC or -PE (STA), CD106 (AF809; R&D Systems), and donkey anti-sheep IgG-Alexa Fluor 488 (Invitrogen) antibodies were used. The relevant isotype antibodies (MOPC-21 and 31243 (Invitrogen) were used as controls.

For intracellular immunostaining of proteins, single-cell suspensions were labeled with LIVE/DEAD Fixable Dead Cell Stain Kit (Green or Aqua; Invitrogen) and then treated with FIX & PERM Fixation & Cell Permeabilization Kit (Invitrogen) according to the manufacturer's methanol modification procedure. Cells were then stained with fluorescently labeled antibodies against anti-mouse CD45.1-PerCP/Cy5.5 (A20), CD45.2-APC/Cy7 (104), CD5-PE/Cy7 (53-7.3), pIGF1R-Alexa Fluor 647 (K74-218; pY1131; BD Biosciences), pAKT-PE or -Brilliant Violet 421 (J24-618; pS473; BD Biosciences), pPYK2-PE (L68-1256.272; pY402; BD Biosciences), pFAK (#3284; pY925; Cell Signaling Technology), donkey anti-rabbit IgG-PE (Poly4064), ILK (P83A9), and goat anti-mouse IgG-Alexa Fluor 488 (Invitrogen) antibodies. The relevant isotype antibodies (MOPC-21, MPC-11, and Poly29108) were used as negative controls.

**In vitro cultures of mouse and human T-ALL cells**

Complete RPMI cell culture media was made up of Roswell Park Memorial Institute 1640 (RPMI 1640) supplemented with 10% fetal bovine serum (FBS; Gemini Bio-Products), 55 μM β-mercaptoethanol, 1× GlutaMAX (2 mM L-alanyl-L-glutamine dipeptide), 1 mM Sodium Pyruvate, 1× MEM Non-essential Amino Acid Solution (Sigma-Aldrich), 1× Penicillin-Streptomycin-Glutamine (100 units of penicillin, 100 μg of streptomycin, and 292 μg/ml of L-glutamine). All reagents were obtained from Gibco, unless otherwise indicated.

For enrichment of mouse T-ALL cells, single-cell suspensions of leukemic spleens from mice engrafted with primary T-ALL were incubated with biotinylated antibodies against anti-mouse CD11b (M1/70), CD11c (N418), F4/80 (BM8), and I-A/I-E (M5/1114.15.2), and negatively enriched with MojoSort Streptavidin Nanobeads, according to the manufacturer's instructions. For preparation of tumor-associated myeloid cells, single-cell suspensions of leukemic spleens were incubated with biotinylated antibodies against CD11c and positively isolated using Streptavidin Microbeads and MACS LS Columns (both from Miltenyi Biotec). $3\times10^5$ T-ALL cells were cultured in the presence or absence of $2\text{-}2.5\times10^4$ enriched myeloid cells in 200 μl of complete RMPI in flat-bottom 96-well plates at 37˚C, 5% $CO_2$ for 6-7 days. All reagents were obtained from BioLegend, unless otherwise indicated.

For culture of human T-ALL cells, deidentified primary pediatric T-ALL samples were obtained from Texas Children's Hospital (Houston, TX). Sample procurement and analysis were approved by the institutional review board committees at The University of Texas at Austin and Texas Children's Hospital/Baylor College of Medicine. Patient T-ALL cells were isolated by density gradient separation using Ficoll, washed, and frozen in RPMI 1640 supplemented with 40% (v/v) FBS and 10% dimethyl sulfoxide (DMSO) upon collection. Human myeloid cells were prepared as previously reported.[16] Briefly, leukoreduction system chambers were obtained from We Are Blood (Austin, TX) and peripheral blood mononuclear cells (PBMCs) were isolated by density gradient separation using Histopaque (1.077 g/ml; Sigma-Aldrich). For enrichment of monocytes, $40\times10^6$ PBMCs were plated in 150×15 mm Petri Dish (Corning) in 25 ml of complete RPMI and incubated at 37˚C, 5% $CO_2$ for 2 hours, after which adherent monocytes were collected using cold PBS and cell scrapers (Fisher Scientific). For enhanced purity, collected monocytes were incubated with biotinylated antibodies against human CD3 (UCHT1; BioLegend) and CD19 (HIB19; BioLegend) and lymphocytes were depleted with MojoSort Streptavidin Nanobeads (BioLegend), according to the manufacturer's instructions. For differentiation of macrophages, $40\times10^6$ PBMCs were seeded into T75 flasks (Corning) in 20 mL of complete RPMI medium and cultured at 37°C, 5% $CO_2$ for 2 hours to allow for monocyte adhesion. After adherent monocytes were washed twice with warm complete RPMI medium, monocytes were differentiated for 7-9 days in the presence of macrophage colony-stimulating factor (M-CSF; 50 ng/mL; PeproTech). M-CSF-derived macrophages were collected using cold PBS and cell scrapers (Fisher Scientific). For co-cultures, $1\times10^5$ patient T-ALL cells were plated alone, with monocytes, or with M-CSF-derived macrophages at a 1:1 or 2:1 ratio, respectively, in 200 µL of complete RPMI in flat-bottom 96-well plates for 6-7 days prior to assessing the viability of T-ALL cells by flow cytometry.

To determine whether blockade of integrin-mediated cell adhesion inhibits myeloid-mediated support of mouse T-ALL, $3\times10^5$ T-ALL cells isolated from primary T-ALL-engrafted mice were co-cultured with $2\text{-}2.5\times10^4$ enriched tumor-associated myeloid cells in the presence or absence of blocking antibodies against integrins (anti-mouse CD11a (M17/4) and/or CD29 (HMβ1-1)) or adhesion molecules (anti-mouse CD54 (YN1/1.7.4) and/or CD106 (429(MVCAM.A)). The relevant isotype antibodies (RTK2758, RTK4530, and HTK888, 53-6.7 (anti-mouse CD8), and 7E.17G9 (anti-mouse ICOS)) were used as controls.

To determine whether inhibition of integrin-mediated cell adhesion suppresses myeloid-mediated support of human T-ALL, $1\times10^5$ T-ALL cells were plated with monocytes or M-CSF-derived macrophages at a 1:1 or 2:1 ratio, respectively, in the presence or absence of a cocktail of anti-human CD54 (HCD54; 20 µg/ml) and CD106 (AF809; R&D Systems) or anti-human CD11a (HI111) and CD29. The relevant isotype antibodies (MOPC-21, 31243 (Invitrogen), and OKT-6 (anti-human CD1a; BioXCell; 30 µg/ml)) were used as controls. The final concentration of each antibody is 10 µg/ml, unless otherwise indicated, and all the antibodies above were purchased from BioLegend, unless otherwise indicated.

To determine whether blockade of integrin-mediated close contact inhibits sensitization of mouse T-ALL cells to exogenous IGF1 by tumor-associated myeloid cells, $3\times10^5$ T-ALL cells were cultured in the presence or absence of $2\text{-}2.5\times10^4$ enriched tumor-associated myeloid cells, with or without blocking antibodies against mouse adhesion molecules (above), and in the presence or absence of 100 ng/ml of recombinant murine IGF1 (Peprotech).

To determine whether inhibition of integrin-mediated close contact diminishes IGF1R signaling, $1\times10^6$ mouse T-ALL cells were co-cultured with $7\text{-}8\times10^4$ myeloid cells in the presence of a cocktail of anti-mouse CD54 (YN1/1.7.4) and CD106 (429(MVCAM.A) antibodies, or the relevant isotype controls, at 10 µg/ml each in 500 µl of complete RPMI in flat-bottom 48-well plates. On day 3-4 of culture, cells in each group were intracellularly stained for phosphorylated proteins as described above.

To determine whether survival of mouse or human T-ALL cells co-cultured with myeloid cells requires focal adhesion signaling (mouse: $3\times10^5$ T-ALL cells were cultured in the presence or absence of 2-$2.5\times10^4$ enriched tumor-associated myeloid cells / human: $1\times10^5$ T-ALL cells were cultured with monocytes or M-CSF-derived macrophages at a 1:1 or 2:1 ratio, respectively), cells were cultured for 4-5 days prior to addition of 0.1-2.5 µM FAK/PYK2 dual inhibitors (PF-431396 and PF-562271; Selleckchem). Viability was assessed by flow cytometry 3-4 days after inhibitor addition in comparison to cells treated with vehicle control (DMSO).

**Transwell assays**

Enriched T-ALL cells and/or myeloid cells from the spleens of mice engrafted with primary LN3 T-ALL were plated in the top or bottom chambers of 12 mm Transwell plates with 0.4 µm pore polyester membrane cell culture inserts (Corning) as indicated in Figure 1A. $1\times10^6$ T-ALL cells or $5\times10^4$ myeloid cells were plated in the top chamber of the inserts, while $3\times10^6$ T-ALL cells +/- $1.5\times10^5$ myeloid cells were plated in bottom chamber, with 500 µl or 1.5 ml of complete RMPI, respectively. The viability of T-ALL cells in each chamber was assessed by flow cytometry after 6-7 days of culture.

**Quantitative PCR**

Thymic T-ALL cells were resuspended in TRIzol (Invitrogen), RNA was extracted, and cDNA was generated using qScript cDNA SuperMix (Quantabio). Real-time PCR experiments were performed on an Applied Biosystems Viia7 instrument with the 7500 Fast real-time PCR system using the following primers: beta-actin forward 5'-CACTGTCGAGTCGCGTCCA-3', beta-actin reverse 5'-CATCCATGGCGAACTGGTGG-3', ITGα4 forward 5'- GCA GAG TCT CCG TCA AGA TTT-3', ITGα4 reverse 5'- CCT GGT GTG TCC TAC ATT TCT C-3', ITGα9 forward 5'- TCC CTG CTA CGA AGA GTA TAA GA-3', ITGα9 reverse 5'- GAG TGT CCC AGC CCA ATA AA-3', ITGαL forward 5'- GCC TAT CCT GAG ACC TTC AAT C-3', ITGαL reverse 5'- AGG TTT GCC TCA CAC TTC TT-3', ITGβ1 forward 5'-GAC AGT GTG TGT GTA GGA AGA G-3', ITGβ1 reverse 5'-GCC TCC ACA AAT TAA GCC ATT AG-3', ITGβ2 forward 5'- GTG GTA GGT GTC GTA CTG ATT G-3', ITGβ2 reverse 5'- GGG ACT TGA GTT TCT CCT TCT C-5'

**Bioinformatic analysis of mouse T-ALL RNA-seq data**

Our published RNA-seq results of LN3 T-ALL vs healthy T-lineage cells were re-analyzed. In brief, FastQ files were assessed for quality using FastQC. Reads were then pseudo-aligned or aligned to the mouse transcriptome (GRCm38) to estimate transcript abundances using Kallisto (1) or HISAT2 (2) respectively. We found they perform similarly, as has been reported (3). The transcript-level counts were aggregated to gene-level counts using tximport in R (4), normalized using DESeq2 (5) and size factors were transformed with variance stabilizing transformation to yield counts that are approximately homoscedastic. Genes that correlated significantly (absolute fold change ≥ 2, adjusted *P*-value < 0.05) to sequencing batch were identified using the likelihood ratio test in DESeq2 and were removed from further analysis. To determine processes upregulated in T-ALL, we compared T-ALL and healthy T-lineage cells across splenic and thymic environments using DESeq2, yielding a list of upregulated genes (fold change ≥ 2, adjusted *P*-value < 0.05), and carried out functional enrichment analyses using Metascape (6) with Gene Ontology (GO) Biological Processes, Kyoto Encyclopedia of Genes and Genomes (KEGG) Pathways, Reactome gene sets, and WikiPathways; default cutoffs were used for enrichment (minimum overlap = 3, *P*-value cutoff = 0.01, minimum enrichment = 1.5). To perform pairwise comparisons of T-ALL cells vs healthy T-lineage cells in the spleens or thymuses separately, Gene Set Enrichment Analysis (GSEA) was performed on differentially expressed genes identified by DESeq2 (absolute fold change ≥ 1.5, adjusted *P*-value < 0.05) using the PID gene sets with gene set permutations (1000) and default parameters (enrichment statistic: weighted, metric for ranking: signal2noise, gene list sorting model: real, gene list ordering mode: descending, maximum size:500, minimum size:15). The top 15 pathways significantly enriched or depleted in T-ALL cells relative to controls in each organ were visualized using the pheatmap package in R (The R Foundation for Statistical Computing). Further pathway analyses using the WikiPathways 2019 Mouse gene sets of Enrichr was run on all genes highly expressed in T-ALL cells in the thymuses relative to thymocytes (fold change ≥ 1.5, adjusted *P*-value < 0.05) (7–9).

**Bioinformatic analysis of patient T-ALL RNA-seq data**

Bioinformatic analysis was performed on published RNA-seq data of T-ALL samples from 264 patients and the associated clinical outcomes datasets from the TARGET ALL Phase II project. Patients whose WBC numbers at diagnosis were higher than 200,000/ml were excluded. We used xCell (10) to quantify the enrichment scores of macrophages and monocytes from bulk RNA-seq data. Pathway information related to integrin signaling was obtained from MSigDB using the R package "msigdbr (11)." The enrichment scores of the integrin pathways in each RNA-seq sample were calculated using the "fgsea" function from the "fgsea" R package (12). For calculation of FAK signaling signature scores, we downloaded the transcriptomics data of FAK-WT and FAK-null murine squamous cell carcinoma cells from Gene Expression Omnibus (GEO; GSE147670), and performed differential expression analyses, identifying the top 50 most up-regulated genes in FAK-WT cells. The 50 mouse genes were mapped to human orthologs using the biomaRt package (13). Using the 50-gene list as an expression signature of FAK signaling, we calculated signature scores for each patient sample. To plot a longitudinal event-free survival of pediatric T-ALL patients, we divided patient data into high and low groups for each integrin or FAK pathway using the median as a cutoff value. Log-rank tests were used to detect the correlation of the integrin or FAK pathways with the event free survival time of T-ALL patients. Pearson correlations were calculated to assess the correlation between each myeloid subset (monocytes or M-CSF-derived macrophages) and integrin or FAK pathways. R was used for the data analyses (14).

**Supplementary Figures**

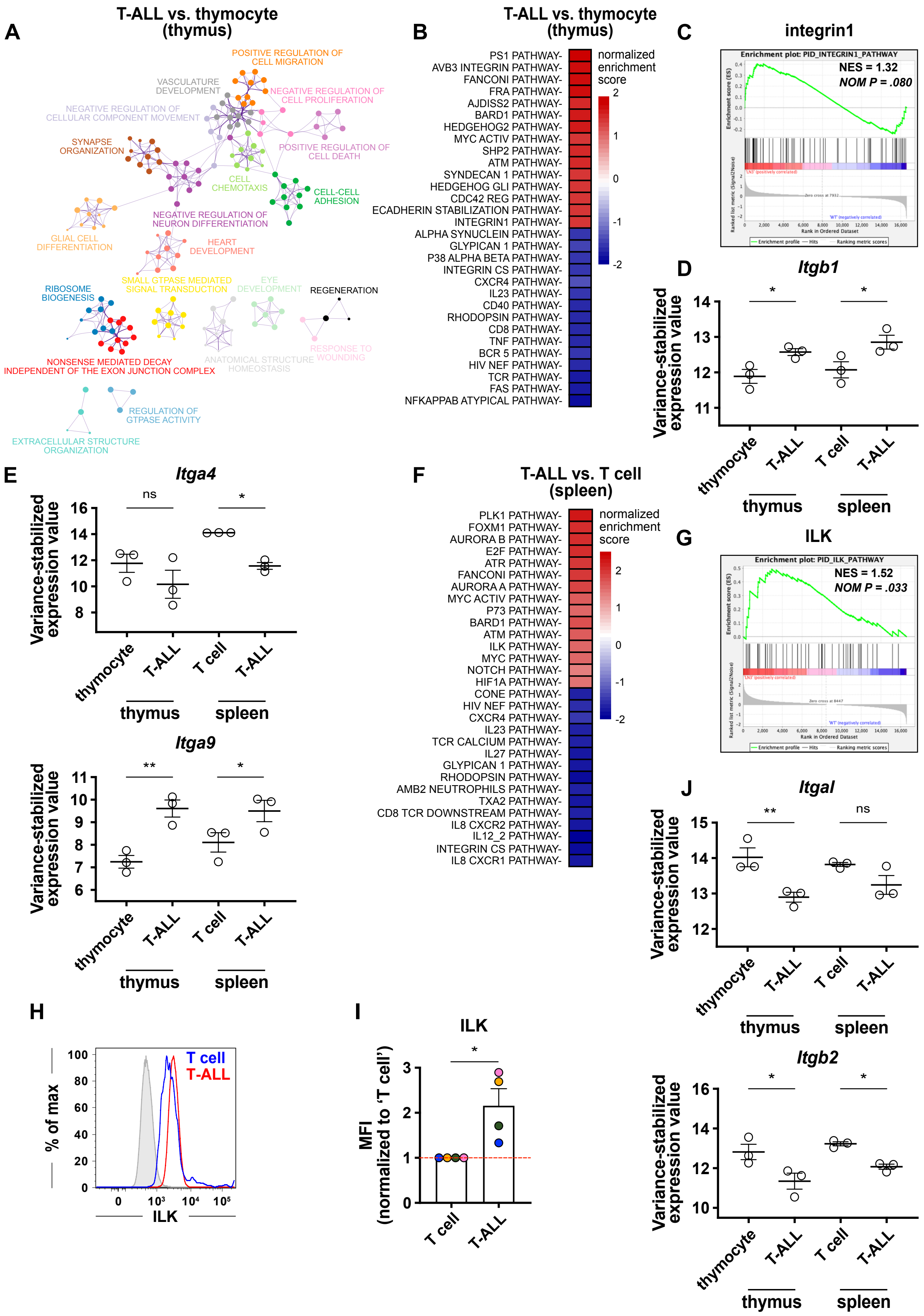
A
T-ALL vs. thymocyte (thymus)
POSITIVE REGULATION OF CELL MIGRATION
VASCULATURE DEVELOPMENT
NEGATIVE REGULATION OF CELL PROLIFERATION
NEGATIVE REGULATION OF CELLULAR COMPONENT MOVEMENT
SYNAPSE ORGANIZATION
POSITIVE REGULATION OF CELL DEATH
CELL CHEMOTAXIS
CELL-CELL ADHESION
NEGATIVE REGULATION OF NEURON DIFFERENTIATION
GLIAL CELL DIFFERENTIATION
HEART DEVELOPMENT
SMALL GTPASE MEDIATED SIGNAL TRANSDUCTION
EYE DEVELOPMENT
REGENERATION
RIBOSOME BIOGENESIS
RESPONSE TO WOUNDING
NONSENSE MEDIATED DECAY INDEPENDENT OF THE EXON JUNCTION COMPLEX
ANATOMICAL STRUCTURE HOMEOSTASIS
REGULATION OF GTPASE ACTIVITY
EXTRACELLULAR STRUCTURE ORGANIZATION
B
T-ALL vs. thymocyte (thymus)
PS1 PATHWAY-
AVB3 INTEGRIN PATHWAY-
FANCONI PATHWAY-
FRA PATHWAY-
AJDISS2 PATHWAY-
BARD1 PATHWAY-
HEDGEHOG2 PATHWAY-
MYC ACTIV PATHWAY-
SHP2 PATHWAY-
ATM PATHWAY-
SYNDECAN 1 PATHWAY-
HEDGEHOG GLI PATHWAY-
CDC42 REG PATHWAY-
ECADHERIN STABILIZATION PATHWAY-
INTEGRIN1 PATHWAY-
ALPHA SYNUCLEIN PATHWAY-
GLYPICAN 1 PATHWAY-
P38 ALPHA BETA PATHWAY-
INTEGRIN CS PATHWAY-
CXCR4 PATHWAY-
IL23 PATHWAY-
CD40 PATHWAY-
RHODOPSIN PATHWAY-
CD8 PATHWAY-
TNF PATHWAY-
BCR 5 PATHWAY-
HIV NEF PATHWAY-
TCR PATHWAY-
FAS PATHWAY-
NFKAPPAB ATYPICAL PATHWAY-
normalized enrichment score
2
1
0
-1
-2
C
integrin1
Enrichment plot: PID_INTEGRIN1_PATHWAY
NES = 1.32
NOM P = .080
D
Itgb1
Variance-stabilized expression value
thymocyte
T-ALL
T cell
T-ALL
thymus
spleen
E
Itga4
ns
Variance-stabilized expression value
thymocyte
T-ALL
T cell
T-ALL
thymus
spleen
Itga9
Variance-stabilized expression value
thymocyte
T-ALL
T cell
T-ALL
thymus
spleen
F
T-ALL vs. T cell (spleen)
PLK1 PATHWAY-
FOXM1 PATHWAY-
AURORA B PATHWAY-
E2F PATHWAY-
ATR PATHWAY-
FANCONI PATHWAY-
AURORA A PATHWAY-
MYC ACTIV PATHWAY-
P73 PATHWAY-
BARD1 PATHWAY-
ATM PATHWAY-
ILK PATHWAY-
MYC PATHWAY-
NOTCH PATHWAY-
HIF1A PATHWAY-
CONE PATHWAY-
HIV NEF PATHWAY-
CXCR4 PATHWAY-
IL23 PATHWAY-
TCR CALCIUM PATHWAY-
IL27 PATHWAY-
GLYPICAN 1 PATHWAY-
RHODOPSIN PATHWAY-
AMB2 NEUTROPHILS PATHWAY-
TXA2 PATHWAY-
CD8 TCR DOWNSTREAM PATHWAY-
IL8 CXCR2 PATHWAY-
IL12_2 PATHWAY-
INTEGRIN CS PATHWAY-
IL8 CXCR1 PATHWAY-
normalized enrichment score
2
1
0
-1
-2
G
ILK
Enrichment plot: PID_ILK_PATHWAY
NES = 1.52
NOM P = .033
J
Itgal
ns
Variance-stabilized expression value
thymocyte
T-ALL
T cell
T-ALL
thymus
spleen
Itgb2
Variance-stabilized expression value
thymocyte
T-ALL
T cell
T-ALL
thymus
spleen
H
T cell
T-ALL
% of max
ILK
I
ILK
MFI (normalized to 'T cell')
T cell
T-ALL

**Supplementary Figure 1. Integrin-associated pathways are enriched in T-ALL cells relative to tumor-free T-lineage cells.**

(A) Visualization of the network of term clusters significantly enriched in thymic T-ALL cells relative to tumor-free thymocytes using the Cytoscape bioinformatics tool (15). Each term is represented by a circle node with size proportional to the number of input genes that fall into that term, and its color represents its cluster identity. (B) The top 15 pathways significantly enriched or depleted in thymic T-ALL cells relative to tumor-free thymocytes were identified by the Gene Set Enrichment Analysis (GSEA) bioinformatics tool using the PID gene sets. (C) GSEA enrichment plots for the INTEGRIN1 pathway in thymic T-ALL cells relative to tumor-free thymocytes as in (B). Normalized enrichment scores (NES) and nominal *P*-values are shown. (D-E) Variance-stabilized expression values of (D) *Itgb1* and (E) *Itga4* and *Itga9* in tumor-free T-lineage and T-ALL cells from the thymus and spleen, as indicated. Bars represent mean +/- SEM; symbols represent individual biologic replicates. (F) The top 15 pathways significantly enriched or depleted in splenic T-ALL cells relative to tumor-free splenic $CD8^+$ T cells were identified by the Gene Set Enrichment Analysis (GSEA) bioinformatics tool using the PID gene sets. (G) GSEA enrichment plots for the ILK pathway in splenic T-ALL cells relative to tumor-free $CD8^+$ T cells as in (F). (H) Representative flow cytometry plots of ILK expression in transplanted LN3 T-ALL cells (red) and host T cells (blue) from the same leukemic spleens. Isotype control stain is shaded in gray. (I) Quantification of the levels of ILK in T-ALL cells ($CD45.2^+CD5^+$) relative to host T cells ($CD45.1^+CD5^+$) from data as in (H), displayed as median fluorescence intensities (MFI). Results were normalized to the MFI of T cells within each experiment. Bars represent the mean + SEM from 4 independent experiments, each with a distinct color-coded primary T-ALL. The red line indicates the normalized mean MFI of T cells. (J) Variance-stabilized expression values of *Itgal* and *Itgb2* in tumor-free T-lineage and T-ALL cells from the thymus and spleen, as indicated. Bars represent mean +/- SEM; symbols represent individual biologic replicates. Statistical significance was determined by (D, E, J) two-way ANOVA with the Holm-Sidak correction, and (I) paired Student t tests; *P*-values: *<0.05, **<0.01. ns, not significant.

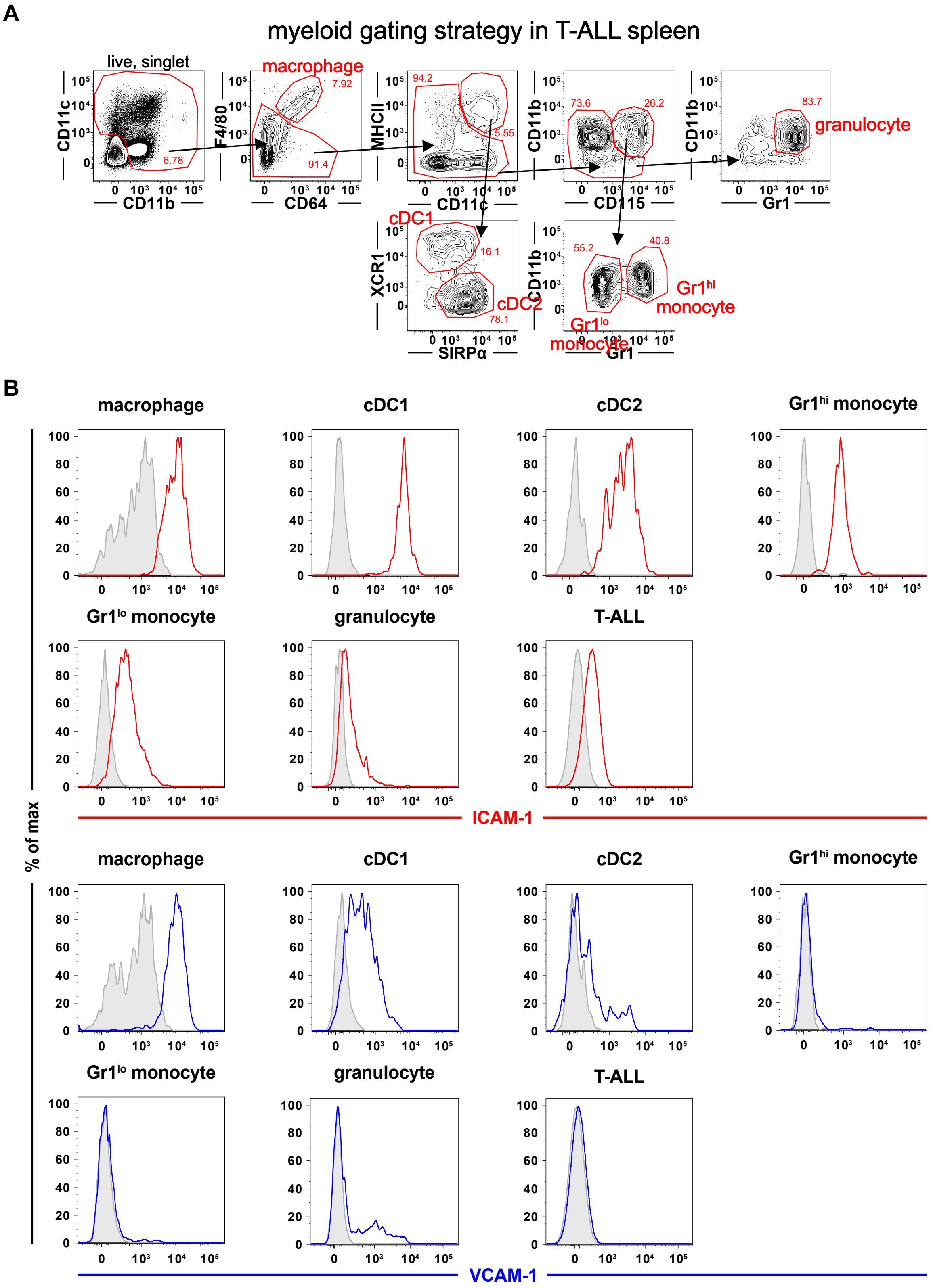
A
myeloid gating strategy in T-ALL spleen
live, singlet
CD11c
CD11b
6.78
macrophage
7.92
F4/80
CD64
91.4
94.2
MHCII
CD11c
5.55
73.6
26.2
CD11b
CD115
83.7
granulocyte
Gr1
cDC1
XCR1
16.1
cDC2
78.1
SIRPα
55.2
40.8
Gr1hi
monocyte
Gr1lo
monocyte
B
macrophage
cDC1
cDC2
Gr1hi monocyte
Gr1lo monocyte
granulocyte
T-ALL
ICAM-1
% of max
VCAM-1

**Supplementary Figure 2. ICAM-1 and VCAM-1 are expressed by tumor-associated myeloid cells.**

(A) Representative sequential flow cytometry gating schemes for evaluation of the indicated myeloid subsets in the spleen of mice transplanted with primary LN3 T-ALL. (B) Representative flow cytometry histograms of ICAM-1 (red) and VCAM-1 (blue) expression by the indicated myeloid subsets or T-ALL cells from spleens of leukemic mice transplanted with primary LN3 T-ALL cells. Isotype control stains are shaded in gray.

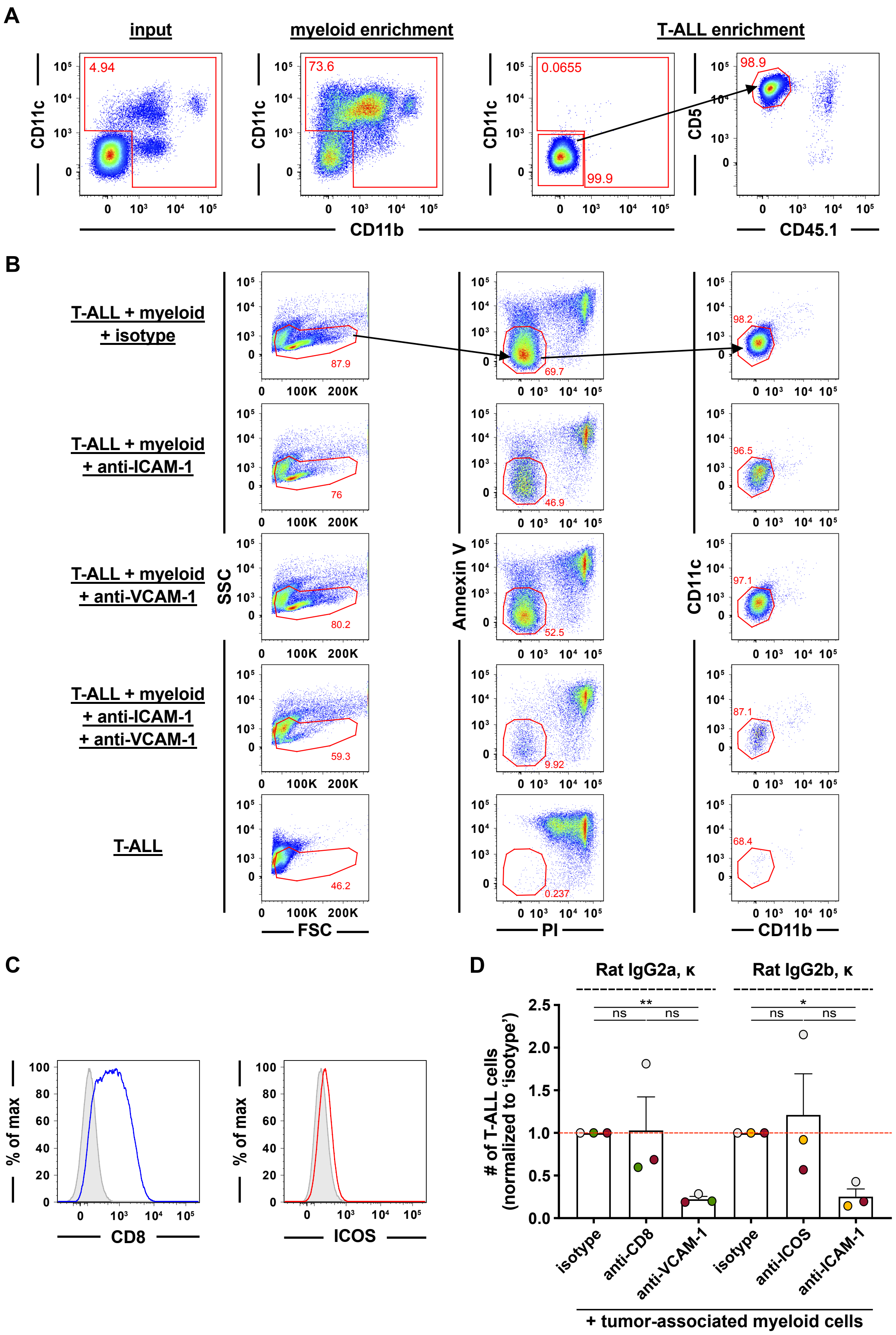
A
input
myeloid enrichment
T-ALL enrichment
CD11c
CD5
CD11b
CD45.1
4.94
73.6
0.0655
99.9
98.9
B
T-ALL + myeloid + isotype
T-ALL + myeloid + anti-ICAM-1
T-ALL + myeloid + anti-VCAM-1
T-ALL + myeloid + anti-ICAM-1 + anti-VCAM-1
T-ALL
SSC
FSC
Annexin V
PI
CD11c
CD11b
87.9
69.7
98.2
76
46.9
96.5
80.2
52.5
97.1
59.3
9.92
87.1
46.2
0.237
68.4
C
% of max
CD8
ICOS
D
Rat IgG2a, κ
Rat IgG2b, κ
### of T-ALL cells (normalized to 'isotype')
ns
**
*
isotype
anti-CD8
anti-VCAM-1
anti-ICOS
anti-ICAM-1
+ tumor-associated myeloid cells

**Supplementary Figure 3. Inhibition of ICAM-1- and VCAM-1-mediated cell adhesion reduces T-ALL survival in vitro.**

(A) Representative flow cytometry plots showing the purity of enriched tumor-associated myeloid ($CD11b^+$ and/or $CD11c^+$) and T-ALL ($CD11b^-CD11c^-CD5^+CD45.2^+$) cells from the spleens of mice transplanted with primary LN3 T-ALL cells with sequential gating indicated by arrows. The “input” column shows myeloid composition ($CD11b^+$ and/or $CD11c^+$) before enrichment. The “myeloid enrichment” column shows the percent of myeloid cells following positive enrichment of $CD11c^+$ cells from the spleens. The third column shows the low frequency of myeloid ($CD11b^-$ and $CD11c^-$) cells within the enriched T-ALL cells following depletion with antibodies against F4/80, CD11b, I-A/I-E, and CD11c. The fourth column shows expression of CD5 and CD45.1 on the enriched T-ALL fraction. (B) Representative flow cytometry plots showing the viability of transplanted LN3 T-ALL cells cultured in the presence or absence of enriched tumor-associated myeloid cells from the spleens, as in (A), and in the presence or absence of anti-ICAM-1 and/or anti-VCAM-1 antibodies. Viable T-ALL cells were quantified as Annexin V $(AV)^-PI^-$ $CD11b^-CD11c^-$ cells. Sequential gating is indicated by arrows. (C) Representative flow cytometry histograms of CD8 (blue) and ICOS (red) expression by T-ALL cells from the spleens of mice transplanted with primary LN3 T-ALL cells. Isotype control stains are shaded in gray. (D) Quantification of viable LN3 T-ALL cells 6-7 days after co-culture with enriched tumor-associated myeloid cells in the presence of anti-CD8 (Rat IgG2a, κ) or anti-ICOS (Rat IgG2b, κ) antibodies (10 μg/ml per each). Data from anti-VCAM-1 (Rat IgG2a, κ)- and anti-ICAM-1 (Rat IgG2b, κ)-treated cultures from Figure 2A were included for comparison. Results were normalized to isotype-treated cultures within each experiment. Bars show the mean + SEM from 3 independent experiments, each with a distinct color-coded primary T-ALL; symbols represent the average of 2-3 technical replicate wells per experiment. The red line indicates the normalized mean viability of isotype-treated T-ALL cells. Statistical significance was determined by (D) repeated measures one-way ANOVA with the Holm-Sidak correction; *P*-values: *<0.05, **<0.01. ns, not significant.

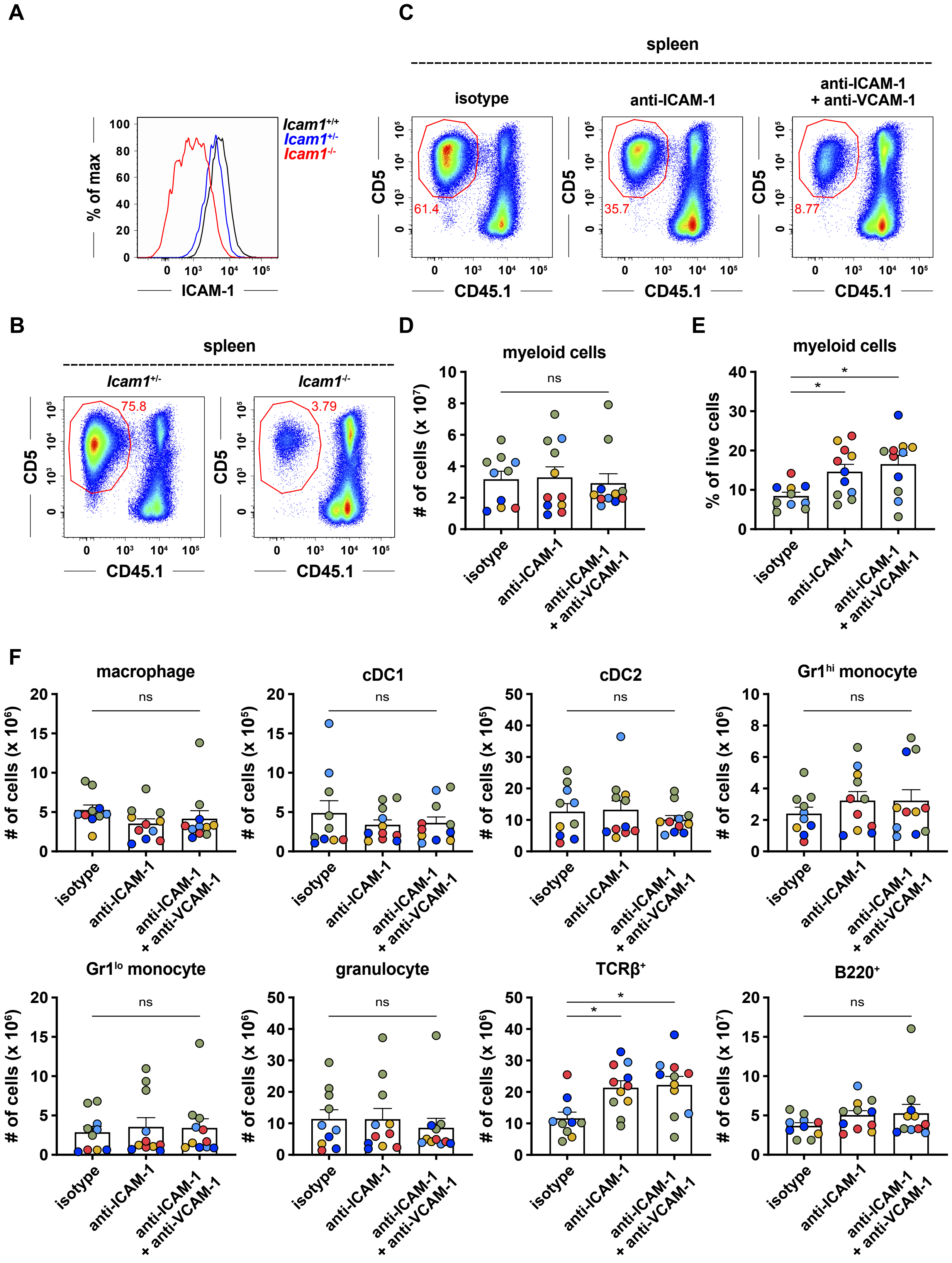
A
% of max
ICAM-1
Icam1+/+
Icam1+/-
Icam1-/-
B
spleen
Icam1+/-
Icam1-/-
75.8
3.79
CD5
CD45.1
C
spleen
isotype
anti-ICAM-1
anti-ICAM-1
+ anti-VCAM-1
61.4
35.7
8.77
CD5
CD45.1
D
myeloid cells
### of cells (x 10^7)
ns
E
myeloid cells
% of live cells
*
*
isotype
anti-ICAM-1
anti-ICAM-1
+ anti-VCAM-1
F
macrophage
cDC1
cDC2
Gr1hi monocyte
Gr1lo monocyte
granulocyte
TCRβ+
B220+
### of cells (x 10^6)
### of cells (x 10^5)
### of cells (x 10^7)
ns

**Supplementary Figure 4. Inhibition of ICAM-1- and VCAM-1-mediated cell adhesion diminishes T-ALL progression in vivo and does not deplete myeloid cells.**
(A) Representative flow cytometry histogram of ICAM-1 expression by macrophages (F4/80$^+$CD64$^+$) from the spleens of *Icam1*$^{+/+}$ (black), *Icam1*$^{+/-}$ (blue), or *Icam1*$^{-/-}$ (red) mice transplanted with primary LN3 T-ALL cells. (B) Representative flow cytometry plots showing a decrease in T-ALL burden in the spleens of LN3 T-ALL-bearing *Icam1*$^{-/-}$ versus *Icam1*$^{+/-}$ mice. (C) Representative flow cytometry plots showing a decrease in T-ALL burden in the spleens of LN3 T-ALL-transplanted mice following treatment with anti-ICAM-1 antibody alone or a cocktail of anti-ICAM-1 and anti-VCAM-1 antibodies (100 µg each per mouse) as in Figure 3F. (D-E) Quantification of the (D) number and (E) frequency of myeloid cells (CD11b$^+$ and/or CD11c$^+$ cells) in the spleens of mice transplanted with primary LN3 T-ALL and treated with either anti-ICAM-1 antibody alone, a cocktail of anti-ICAM-1 and anti-VCAM-1 antibodies, or isotype control antibodies, as in (C). (F) Quantification of the indicated myeloid subsets or TCRβ$^+$ or B220$^+$ host cells in the spleens of mice from the same experiments as in (D-E). Statistical significance was determined by (D, E, F) repeated measures one-way ANOVA with the Holm-Sidak correction; *P*-values: *<0.05. ns, not significant.

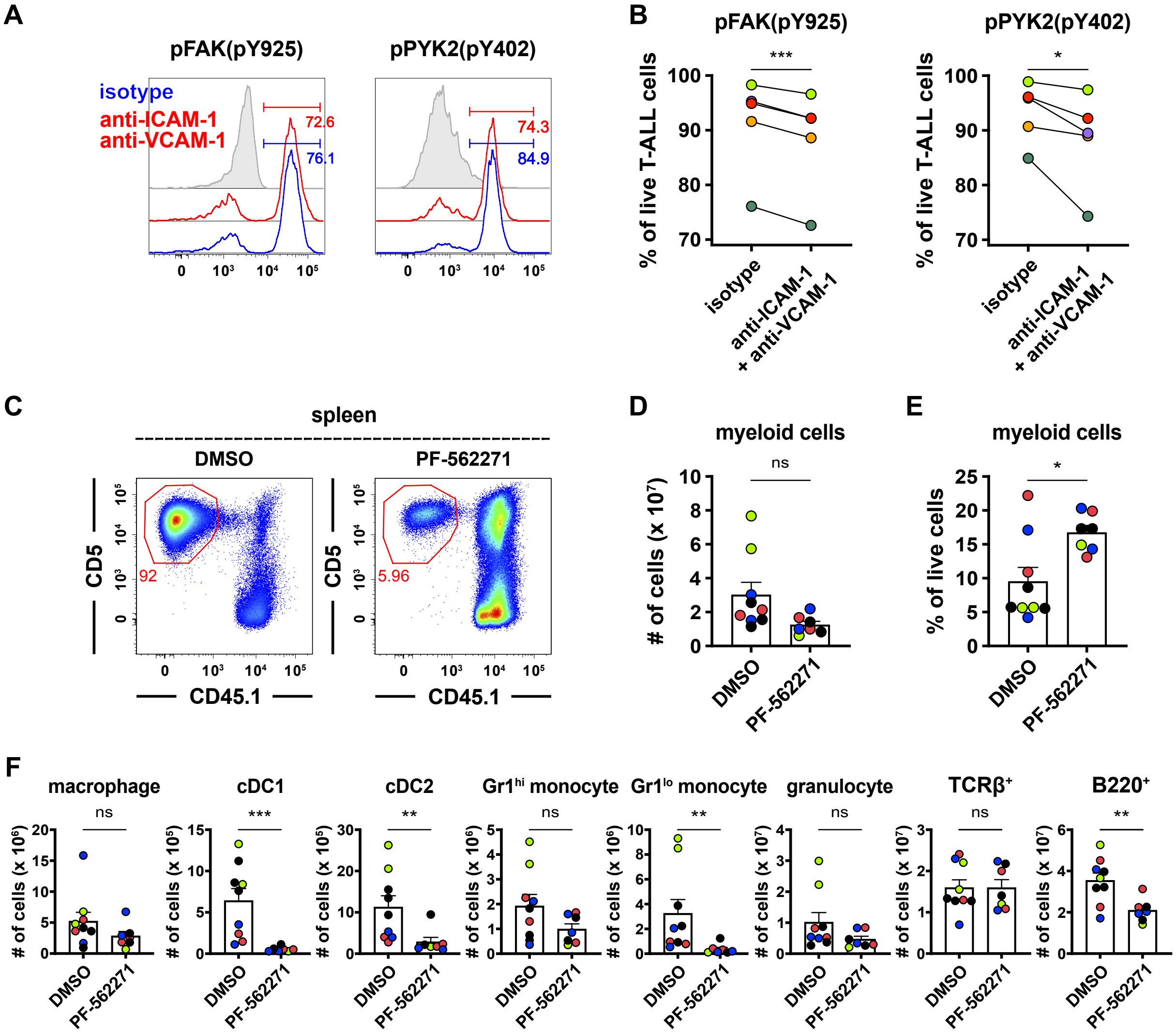
A
pFAK(pY925)
pPYK2(pY402)
isotype
anti-ICAM-1
anti-VCAM-1
72.6
76.1
74.3
84.9
B
% of live T-ALL cells
isotype
anti-ICAM-1
+ anti-VCAM-1
***
*
C
spleen
DMSO
PF-562271
CD5
CD45.1
92
5.96
D
myeloid cells
### of cells (x 10^7)
ns
E
myeloid cells
% of live cells
*
F
macrophage
cDC1
cDC2
Gr1^hi monocyte
Gr1^lo monocyte
granulocyte
TCRβ+
B220+
### of cells
ns
***
**
ns
**
ns
ns
**
DMSO
PF-562271

**Supplementary Figure 5. Inhibition of ICAM-1- and VCAM-1-mediated adhesion diminishes FAK and PYK2 signaling in T-ALL cells and reduces leukemia burden in vivo.**
(A) Representative flow cytometry histograms of pFAK (left) and pPYK2 (right) expression in LN3 T-ALL cells 3-4 days after co-culture with tumor-associated myeloid cells in the presence of blocking antibodies against ICAM-1 and VCAM-1 (red) or isotype controls (blue). Isotype control stains are shaded in gray. (B) Quantification of the frequency of viable T-ALL cells expressing $pFAK^+$ or $pPYK2^+$, as indicated, from experiments as in (A). (C) Representative flow cytometry plots showing a decrease in T-ALL burden in the spleens of LN3 T-ALL-transplanted mice treated as per Figure 4E with PF-562271 (a FAK/PYK2 dual inhibitor; 25 mg per kg body weight) or DMSO. (D-E) Quantification of the (D) number and (E) frequency of myeloid cells ($CD11b^+$ and/or $CD11c^+$ cells) in the spleens of LN3 leukemic mice following treatment with either PF-562271 or DMSO, as in (C). (F) Quantification of the indicated splenic myeloid subsets or $TCR\beta^+$ or $B220^+$ host cells from the same experiments as in (D-E). Data in (C-F) are from the same experiments as in Figure 4E-G. Statistical significance was determined by (B) paired Student t tests, and (D, E, F) unpaired Student t tests; *P*-values: *<0.05, **<0.01, ***<0.001. ns, not significant.

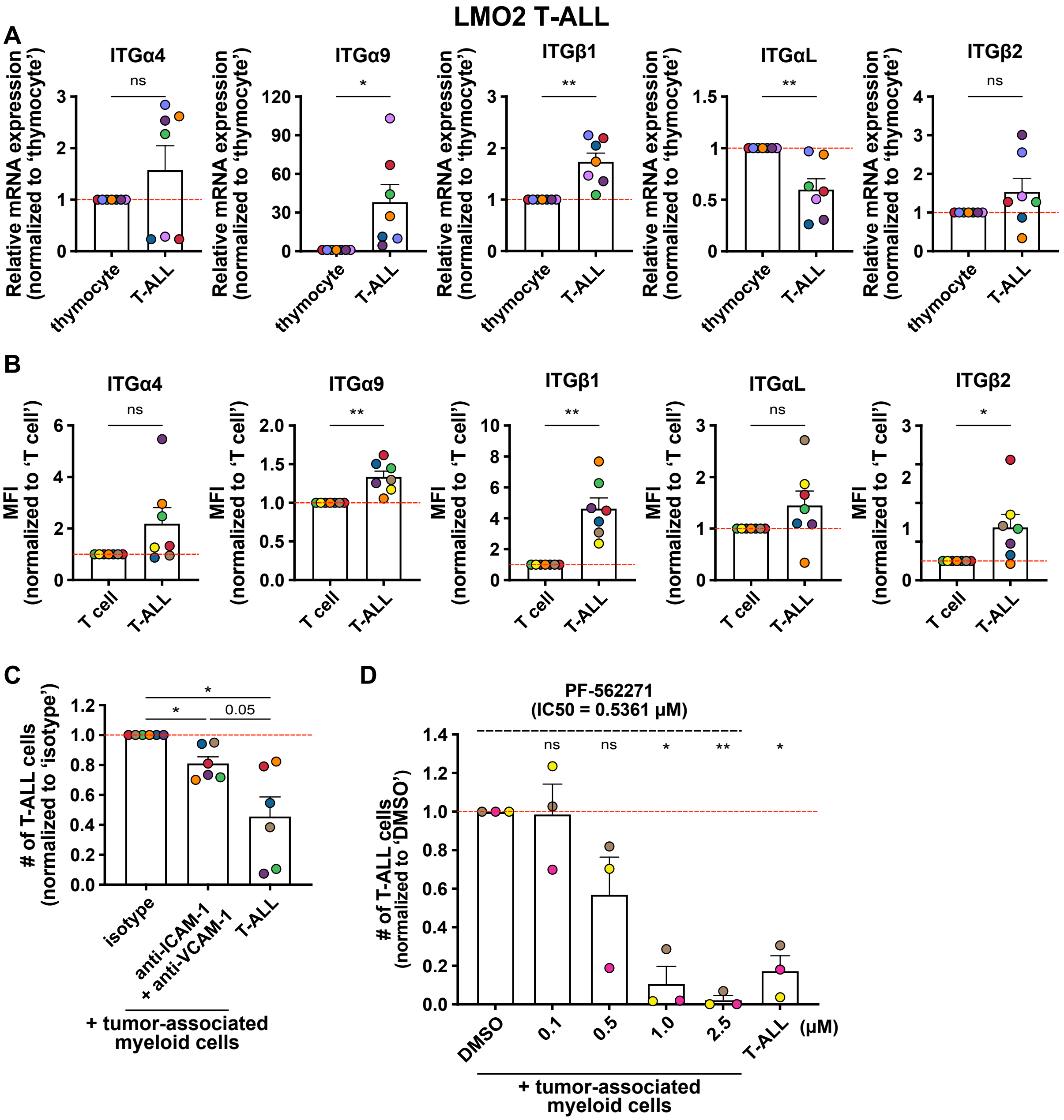
LMO2 T-ALL
A
ITGα4
ITGα9
ITGβ1
ITGαL
ITGβ2
Relative mRNA expression (normalized to 'thymocyte')
thymocyte
T-ALL
ns
*
**
B
MFI (normalized to 'T cell')
T cell
C
### of T-ALL cells (normalized to 'isotype')
isotype
anti-ICAM-1 + anti-VCAM-1
0.05
+ tumor-associated myeloid cells
D
PF-562271 (IC50 = 0.5361 μM)
### of T-ALL cells (normalized to 'DMSO')
DMSO
(μM)

**Supplementary Figure 6. Tumor-associated myeloid cells support T-ALL survival in an integrin-dependent manner in the LMO2 mouse of T-ALL.**

(A) Relative transcript levels of the indicated integrin subunits were quantified by qRT-PCR in healthy thymocytes and in primary thymic LMO2 T-ALL cells. Results were normalized to the expression level of tumor-free thymocytes within each experiment. Bars represent mean + SEM from 7 independent experiments, each with a distinct color-coded primary T-ALL. The red line indicates the normalized mean levels of tumor-free thymocytes. (B) Protein expression of the indicated integrin subunits was quantified by flow cytometry on transplanted LMO2 T-ALL cells (CD45.2$^+$CD5$^+$) and host T cells (CD45.1$^+$CD5$^+$) from leukemic spleens. MFI intensity values were normalized to the MFI of host T cells within each experiment. Data are compiled from 7 independent experiments, each with a distinct color-coded primary LMO2 T-ALL. (C) Quantification of viable splenic LMO2 T-ALL cells was assessed 6-7 days after culture alone or in the presence of enriched tumor-associated myeloid cells and anti-ICAM-1/anti-VCAM-1 antibodies (10 µg/ml per each) or isotype controls, as indicated. Results were normalized to isotype-treated cultures. Bars show the mean + SEM from 6 independent experiments, each with a distinct color-coded primary T-ALL; symbols represent the average of 2-3 technical replicates per experiment. The red line indicates the normalized mean viability of isotype-treated T-ALL cells. (D) Quantification of viable LMO2 T-ALL cells co-cultured with enriched tumor-associated myeloid cells for 4 days before addition of the indicated concentrations of a FAK/PYK2 dual inhibitor (PF-562271) or vehicle control (DMSO). Viability was assessed after 2-3 days. Results were normalized to DMSO-treated cultures in each experiment. Bars represent means + SEM from 3 independent experiments using distinct color-coded primary T-ALLs; symbols represent the average of 2-3 technical replicate wells. The red line indicates the normalized mean T-ALL viability in DMSO-treated cultures. Statistical significance was determined by (A, B) paired Student t tests, and (C, D) repeated measures one-way ANOVA with the Holm-Sidak correction; *P*-values: *<0.05, **<0.01. ns, not significant.

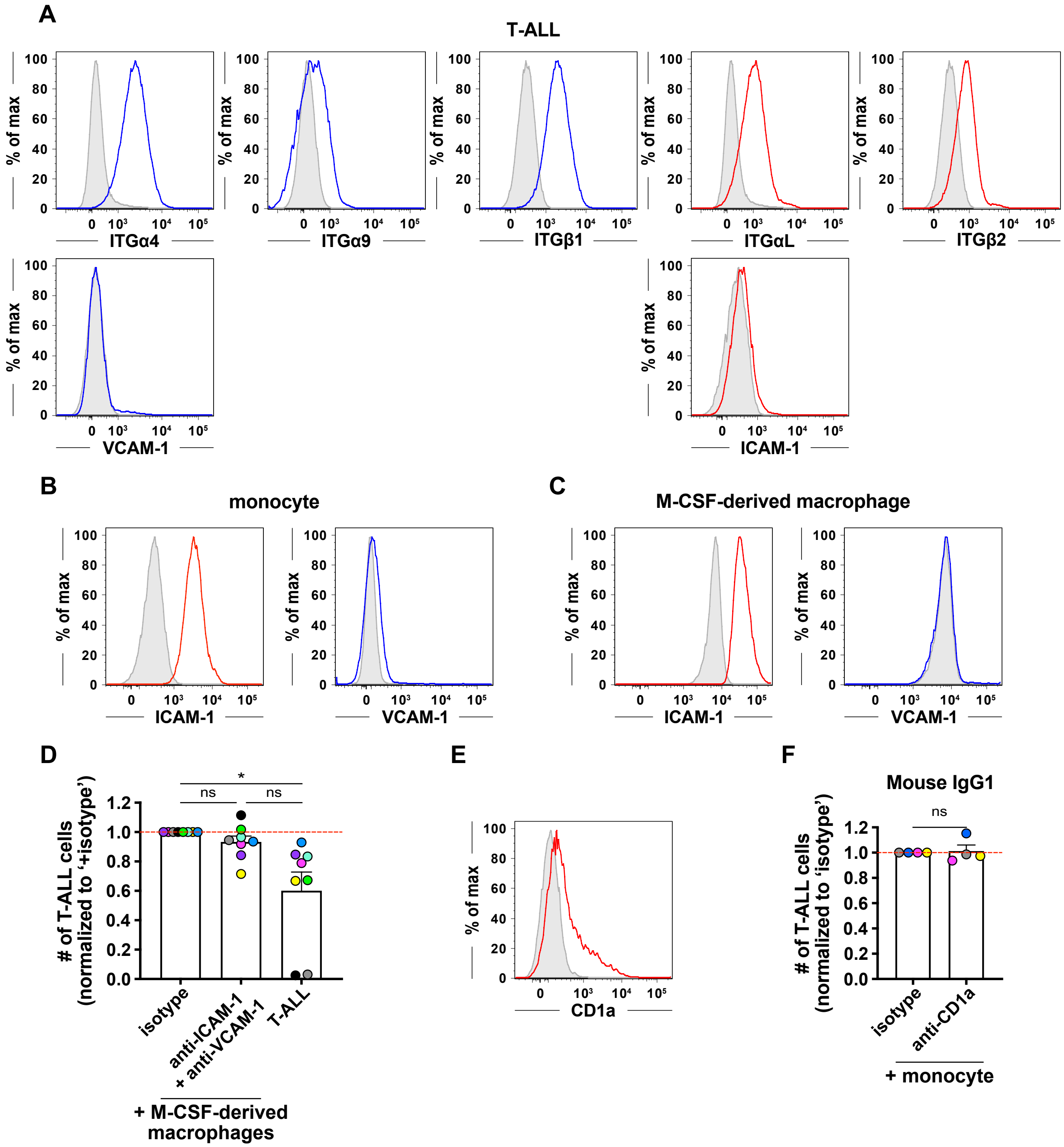
A
T-ALL
% of max
ITGα4
ITGα9
ITGβ1
ITGαL
ITGβ2
VCAM-1
ICAM-1
B
monocyte
C
M-CSF-derived macrophage
D
### of T-ALL cells (normalized to '+isotype')
ns
*
isotype
anti-ICAM-1 + anti-VCAM-1
T-ALL
+ M-CSF-derived macrophages
E
CD1a
F
Mouse IgG1
### of T-ALL cells (normalized to 'isotype')
anti-CD1a
+ monocyte

**Supplementary Figure 7. Integrins and adhesion molecules are expressed by primary patient T-ALL cells and human myeloid cells, respectively, and enriched gene signatures of the FAK pathway correlate with inferior patient outcomes.**
(A) Representative flow cytometry histograms of the indicated integrin components or adhesion molecules present on primary patient T-ALL cells. Integrin components binding to VCAM-1 and ICAM-1 are shown in blue and red, respectively. Isotype control stains are shaded in gray. (B-C) Representative flow cytometry histograms showing ICAM-1 (red) and VCAM-1 (blue) expression by (B) PBMC-derived monocytes and (C) M-CSF-derived macrophages. Isotype control stains are shaded in gray. (D) Quantification of viable primary patient T-ALL cells cultured for 6-7 days alone or with M-CSF-derived macrophages. Co-cultures were carried out in the presence of (A) anti-ICAM-1 (20 µg/ml) and anti-VCAM-1 (10 µg/ml) blocking antibodies or isotype control antibodies as indicated. Results were normalized to isotype-treated cultures in each experiment. Bars represent means + SEM from 7 independent experiments using 8 distinct, color-coded patient-derived T-ALLs; symbols represent the average of 2 technical replicate wells. The red line indicates the normalized mean T-ALL viability in isotype-treated cultures. (E) Representative flow cytometry plots of CD1a expression by primary patient T-ALL cells. Isotype control stain is shaded in gray. (F) Quantification of viable patient T-ALL cells 6-7 days after co-culture with PBMC-derived monocytes in the presence of anti-CD1a (Mouse IgG1; 30 µg/ml). Results were normalized to isotype-treated cultures in each experiment. Bars represent means + SEM from 4 independent experiments using distinct, color-coded patient-derived T-ALLs; symbols represent the average of 2-3 technical replicate wells. The red line indicates the normalized mean T-ALL viability in isotype-treated cultures. Statistical significance was determined by (D) repeated measures one-way ANOVA with the Holm-Sidak correction, (F) paired Student t tests. *P*-values: *<0.05. ns, not significant.
